# Mitochondrial Signatures Shape Phenotype Switching and Apoptosis in Response to PLK1 and RSK Inhibitors in Melanoma

**DOI:** 10.1101/2024.06.14.599035

**Authors:** Émilie Lavallée, Maëline Roulet-Matton, Viviane Giang, Roxana Cardona Hurtado, Dominic Chaput, Simon-Pierre Gravel

## Abstract

PLK1 inhibitors are emerging anti-cancer agents being tested in monotherapy and combination therapies for various cancers. Although PLK1 inhibition in experimental models shows potent antitumor effects, translation to the clinic has been hampered by low antitumor activity and tumor relapse. Here, we report the identification of mitochondrial protein signatures that determine sensitivity to approaches targeting PLK1 in human melanoma cell lines. In response to PLK1 inhibition or gene silencing, resistant cells adopt a pro-inflammatory and dedifferentiated phenotype, while sensitive cells engage apoptosis. Mitochondrial DNA depletion and silencing of the ABCD1 transporter sensitize cells to PLK1 inhibition and attenuate the associated pro-inflammatory response. We also found that non-selective inhibitors of the p90 ribosomal S6 kinase (RSK) exert their anti-proliferative and pro-inflammatory effects via PLK1 inhibition. This work reveals overlooked impacts of PLK1 on phenotype switching and suggests that mitochondrial precision medicine can help improve response to targeted therapies.

## INTRODUCTION

Melanoma is a skin cancer originating from melanocytes, which are specialized epidermis cells responsible for melanin deposition and protection from UV rays. While early lesions are easily managed, late-stage cancer poses significant challenges for treatment due to its aggressive nature and propensity to metastasize. Melanoma is responsible for over 80 % of skin cancer deaths, and the 5-year survival rate of advanced melanoma (stage IV) melanoma is less than 30 % ^1^. Mutations in the BRAF kinase are observed in more than 50 % of melanoma patients, and other frequent melanoma subtypes involve activating mutation of RAS isoforms (20-30 % of patients) or loss of NF-1 (10-15 %) ^2,3^. BRAF mutations lead to constitutive activation of the pro-tumorigenic mitogen-activated protein kinase cascade (BRAF-MEK-ERK) ^4,5^. Despite the improvement of progression-free survival in patients treated with BRAF inhibitors in combination with MEK inhibitors, the emergence of drug resistance through a variety of mechanisms remains a major obstacle in implementing sustainable responses and improving patient outcomes.

The p90 ribosomal S6 kinase (RSK) family contains four members, RSK1-4, that share around 80 % homology. RSK isoforms have context-specific expression patterns and exert diverse functions by phosphorylating many protein substrates (review in ^6^). Specifically, RSK1/2 were shown to be up-regulated in melanoma cells and to act as predominant targets of ERK in melanoma ^7–9^. RSK1/2 were also shown to mediate resistance to chemotherapy and BRAF inhibition ^7,10^. Recent reports have proposed that RSK1/2 are potential therapeutic target in melanoma with constitutive MAPK activation to overcome the resistance to BRAF and MEK inhibitors ^11,12^. Despite the importance of these findings and the identification of new drug candidates, it is crucial to consider the selectivity of RSK inhibitors since some off-target effects may contribute to the observed anti-tumoral activity. One notorious unselective RSK inhibitor is BI-1870, which was shown to bind the active site of dozens of unrelated kinases such as Polo-like kinase (PLK) 1/3, aurora kinase B, PIK3, and exert RSK-independent effects (^13–16^ and reviewed in ^17,18^). For other promising compounds with good biodistribution, tolerability and specific anti-tumoral activities, data on selectivity is often incomplete or unpublished, which may be detrimental to the identification of molecular targets and understanding the mechanism of drug action. Thus, we hypothesized that the anti-proliferative effects of several RSK inhibitors might be caused by off-target inhibition of other protein kinases, such as PLK1, a master regulator of cell division and candidate for targeted therapy with increased expression in melanoma ^19–21^. This possibility prompted us to revisit the real impact of selective RSK inhibition on melanoma proliferation and answer outstanding questions such as effects on tumor inflammation and phenotype switching. Our systematic investigation of RSK inhibitors with distinct specificity profiles allowed us to decipher RSK-specific roles in inflammation and antigen presentation. Our results show that PLK1 is a putative target of unselective RSK inhibitors and mediates their anti-proliferative and pro-inflammatory effects. In addition, they indicate that the marked heterogeneity of the response to RSK and PLK1 inhibition is shaped by the expression of mitochondrial proteins associated with a mitochondrial resistance signature.

## RESULTS

### The anti-proliferative and pro-inflammatory effects of RSK inhibitors in melanoma cells are inhibitor-and cell line-specific

To first examine the association between RSK expression and inflammation in human melanoma, we performed correlation analyses using the Skin Cutaneous Melanoma (SKCM) RNA-Seq data set of human melanoma tumors from TCGA PanCancer Atlas. We also included the analysis of transcripts encoding for putative off-targets of BI-D1870, a notorious non-selective RSK inhibitor (reviewed in ^18^). These analyses revealed that all RSK family members correlate positively with inflammation-associated transcripts (**Figure 1A**), while several BI-D1870 targets, such as PLK1, CDK2, and PTK2, showed a 4 negative correlation. Next, the proliferative impact of knockout and gene silencing of the same kinases on melanoma cell lines was examined using the DepMap portal. Knockout and gene silencing of RSK family members had an overall minimal or inexistent effect on the proliferation of melanoma cell lines, while the targeting of PLK1 had a dramatic effect, followed by the targeting of Aurora B, FAK, and CDK2 (**Figure 1B** and **S1A**). These analyses suggest that the anti-proliferative and pro-inflammatory effects of RSK inhibitors such as BI-D1870, BRD7389, PMD-026, AF007, and fisetin in previous melanoma studies ^11,12,22–25^ could be due to the unselective targeting of other kinases, such as PLK1 (**Figure S1B**).

**Figure 1.**
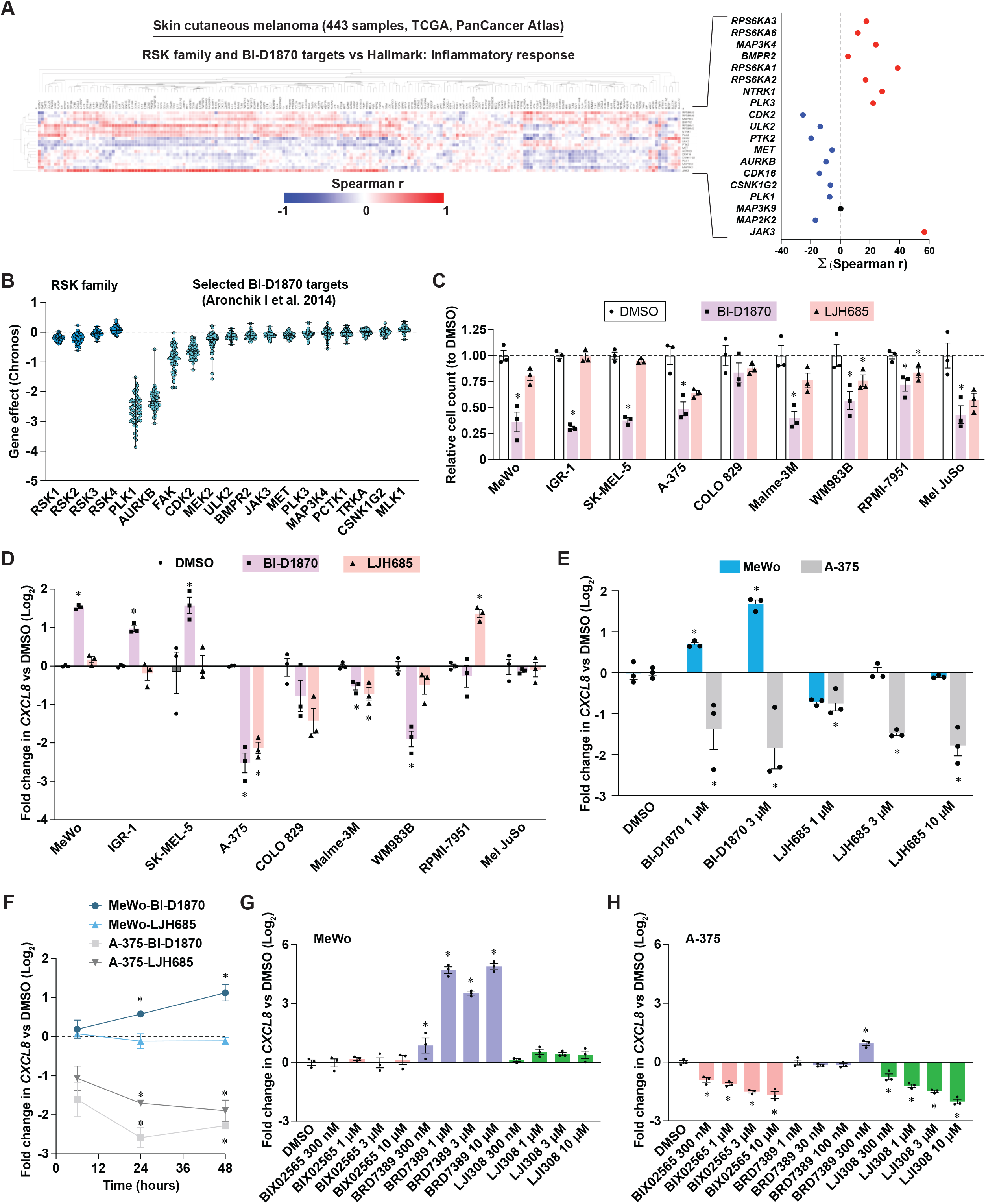
The anti-proliferative and pro-inflammatory effects of RSK inhibitors in melanoma cells are inhibitor-and cell line-specific. See also Figure S1. Correlation of transcripts encoding for RSK family members and BI-D1870 putative targets with inflammatory response transcripts (Hallmarks, MSigDB). The right graph plots the summation of all Spearman correlations in the heat map on the left. (A) Impact of CRISPR loss of function of genes shown in A on proliferation dynamics (Chronos algorithm) of human melanoma cell lines (DepMap). The red line (-1) is the median of all common essential gene scores. Each dot represents a single cell line; data are shown as median ± min/max. (B) Impact of RSK inhibitors on the proliferation of human melanoma cell lines. Cells were treated for 72 h with DMSO 0.1 % (v/v) or inhibitors at 3 µM. (C) *CXCL8* transcript levels in human melanoma cell lines treated as in C. (D) *CXCL8* transcript levels in human melanoma cell lines treated as in C but with escalating doses of RSK inhibitors. (E) *CXCL8* transcript levels in human melanoma cell lines treated as in C but for 6, 24 and 48 h. (G-H) *CXCL8* transcript levels in MeWo and A-375 cell lines in response to escalating doses of additional RSK inhibitors. Data are shown as the means of 3 independent experiments ± SEM. * p < 0.05, One-Way ANOVA with Dunnett multiple comparison test.

To examine the anti-proliferative and pro-inflammatory effects of two RSK inhibitors, the unselective BI-D1870 ^16^ and the selective LJH685 ^13^, we used a panel of nine human melanoma cell lines. BI-D1870 had significant anti-proliferative effects on 8/9 cell lines, while LJH685 had little to no effects (**Figure 1C**). To study the impact of these inhibitors on inflammation, we measured the transcript levels of interleukin 8 (IL-8, gene *CXCL8*), a chemokine linked to melanoma progression, phenotype switching and resistance (reviewed in ^26^). Strikingly, the response to BI-D1870 and LJH685 appeared to be compound-and cell line-specific (**Figure 1D**). BI-D1870 had a pro-inflammatory effect in three melanoma cell lines, MeWo, SK-MEL-5, and IGR-1, while LJH685 had no effect in these cell lines or even had anti-inflammatory effects in A-375, COLO 829, Malme-3M, and WM983B cells. Given their clear opposite response to inhibitors, we selected MeWo and A-375 cell lines for subsequent experiments. At the dose used in the panel (3 µM), the impact of drugs on viability was minimal in both cell lines (**Figure S1C and S1D**). We also observed a dose-dependent effect of drugs on the induction and repression of *CXCL8* in MeWo and A-375 cells, respectively (**Figure 1E**). Since *CXCL8* levels were measured 72 h after stimulation, we asked whether the anti-inflammatory effect of RSK inhibitors was due to negative feedback. In cells treated for shorter periods (6, 24, and 48 h), RSK inhibition with BI-D1870 and LJH685 showed reduced level of *CXCL8* levels in A-375 cells after 24 h, while in MeWo cells an increase in *CXCL8* was observed at the same time point with BI-D1870 (**Figure 1F**). To further 5 strengthen these observations, we examined the impact of additional RSK inhibitors on *CXCL8* in MeWo and A-375 cells. Treatment with the unselective RSK inhibitor BRD7389 ^27^ also strongly induced *CXCL8* in MeWo cells (**Figure 1G**). Interestingly, the pro-inflammatory effects of BRD7389 could not be observed in A-375 above 300 nM due to cell death (**Figure 1H**). Two additional selective RSK inhibitors, BIX02565 ^16^ and LJI308 ^13^, had no effect in MeWo cells but had strong dose-dependent anti-inflammatory effects in A-375 cells (**Figure 1G and 1H**). Taken together, these results reveal an important heterogeneity of response to RSK inhibitors and that other targets than RSK, such as PLK1, may be responsible for the induction a pro-inflammatory response or apoptosis in a cell type-dependent manner.

### Non-selective RSK inhibition induces a pro-inflammatory gene expression program linked to phenotype switching

To further characterize the anti-proliferative and pro-inflammatory effects of the non-selective RSK inhibitor BI-D1870, we performed RNA-Seq analyses in MeWo cells treated for three days with this compound. As expected, a detailed proliferation analysis of MeWo cells shows that BI-D1870 leads to a rapid and sustained proliferation arrest (**Figure 2A**). Next, we performed gene functional classification of differentially expressed transcripts (**Figure 2B**) using the Reactome collection of gene sets on g:Profiler. Up-regulated transcripts were linked to pathways such as neutrophil degranulation, innate immune system, and cellular stress responses (**Figure 2C** and **S2A**), while down-regulated transcripts were almost exclusively linked to cell cycle pathways (**Figure 2D** and **S2B**). Gene set enrichment analyses (GSEA) of normalized reads using the Hallmarks collection of gene sets confirmed the induction of an inflammatory response and the repression of cell cycle genes linked to E2F and checkpoints (**Figure 2E** and **S2C**). Interestingly, Cytoscape network representation of GSEA using the GO:BP collection of gene sets reveals a strong clustering of genes within an immune response cluster that evokes cytoskeletal changes and cell migration (**Figure 2F**). Since the inflammation marker *CXCL8* was among the most enriched up-regulated transcripts from the Inflammatory Response gene set (**Figure 2G**), we next asked whether the corresponding cytokine IL-8 was secreted by MeWo after treatment with BI-D1870. The analysis of cytokines from conditioned media of MeWo revealed a 7-fold induction in IL-8 (**Figure 2H and 2I**) as well as significant induction of other cytokines associated with melanoma progression and therapeutic resistance, such as GRO*α* (*CXCL1*) ^28,29^ and IL-6 ^30^ (**Figure S2D and S2E**). Since the proliferative and inflammatory statuses are tightly associated with phenotype switching in melanoma (reviewed in ^31^), we examined the impact of BI-D1870 on the expression of transcripts previously associated with seven phenotypic states (from differentiated/melanocytic to undifferentiated) ^32^. Strikingly, BI-D1870 had a complex impact on these transcripts, leading to a polarization towards both the melanocytic stage and neural crest-like/undifferentiated states and a depletion of transitory stages (**Figure 2J**). Treatment of MeWo cells with BI-D1870 reduced the expression of Dopachrome Tautomerase/TRP-2 (*DCT*), a modulator of the melanocytic state ^33,34^, while treatment with LJH685 had no impact (**Figure 2K and 2L**). Taken together, these results suggest that non-selective RSK inhibitors such as BI-D1870 can influence melanoma phenotype switching and the tumor immune landscape.

**Figure 2.**
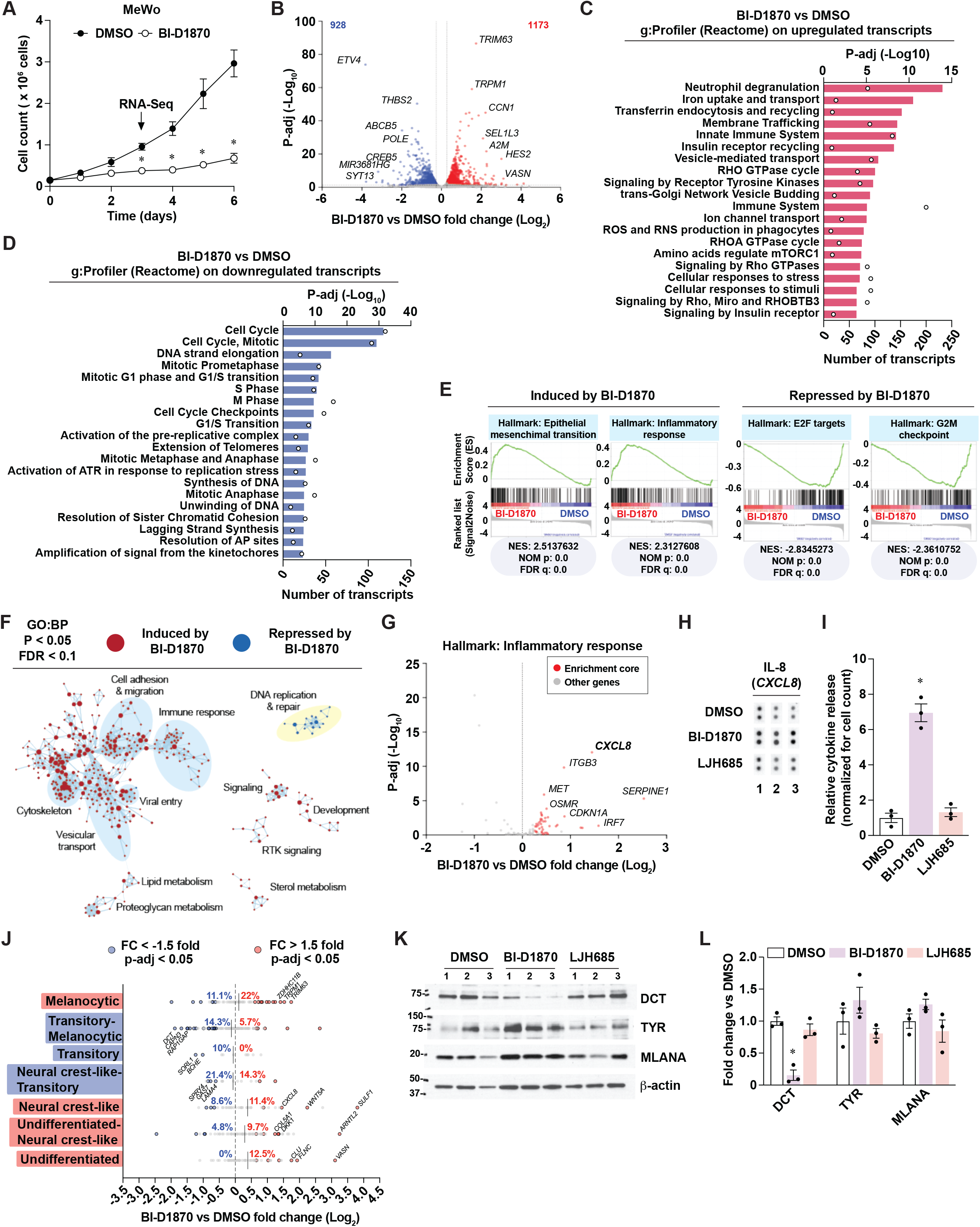
The unselective RSK inhibitor BI-D1870 induces a pro-inflammatory response linked to phenotype switching. See also Figure S2. (A) Proliferation curve in MeWo cells treated with DMSO 0.1 % (v/v) or BI-D1870 (3 µM) for 72 h. (B) Volcano plot that indicates differentially expressed transcripts in MeWo cells treated as in A. Numbers in upper corners indicate the number of up-regulated (red) and down-regulated (blue) significant transcripts based on threshold values of fold change > 1.2 and p-adj < 0.05. (C) Functional gene analysis (g:Profiler; Reactome) of significantly up-regulated transcripts in B. (D) Functional gene analysis as in C for significantly down-regulated transcripts in B. (E) GSEA on selected significantly enriched gene sets (Hallmarks, MSigDB) on normalized data linked to panel B. NES: normalized enrichment score; NOM p: nominal p-value; FDR: False-discovery rate q-value. (F) Visualization of significantly enriched gene sets linked to panel B with Cytoscape (GO:BP, MSigDB, with p < 0.05 and FDR < 0.1). (G) Volcano plot of the inflammatory response gene set (Hallmarks, MSigDB), as indicated in B. Enrichment core (red dots) indicates genes from the GSEA leading edge. (H) IL-8 detection by cytokine array in conditioned media from MeWo cells treated with DMSO 0.1 % (v/v), BI-D1870 (3 µM), or LJH685 (3 µM) for 72h. Numbers indicate independent experiments. (I) Densitometric analysis of 3 independent cytokine arrays related to H. (J) Modulation by BI-D1870 of transcripts associated with phenotype switching in melanoma, done on RNA-Seq data related to B. Percentages indicate the proportion of significantly up-regulated or down-regulated transcripts in each gene set associated with a specific differentiation stage. (K) Immunoblotting of differentiation markers in cell extracts from MeWo cells treated as in H. Numbers above pictures indicate paired independent experiments. (L) Densitometric analysis of immunoblotting from 3 independent experiments related to K. All data are shown as the means of 3 independent experiments ± SEM. A: * p < 0.05, paired Student’s *t*-test. I, L: * p < 0.05, One-Way ANOVA with Dunnett multiple comparison test.

### Selective RSK inhibition is associated with an anti-inflammatory gene expression program and immunometabolic rewiring

To examine the anti-inflammatory effects of selective RSK inhibition, we performed RNA-Seq analyses on A-375 cells treated with LJH685. Treated cells proliferated as control ones at 72 h time-point as well as over a 6-day period (**Figure 3A**). Noticeably, the pro-inflammatory transcript *CXCL8* was among the top differentially expressed transcripts in A-375 cells (**Figure 3B**). Functional gene classification of differentially expressed transcripts in these cells revealed the enrichment of diverse pathways linked to extracellular matrix remodeling, signaling, and metabolism (**Figure 3C and 3D**). Further characterization of down-regulated metabolic transcripts allowed us to identify 197 transcripts implicated in diverse metabolic pathways, such as the metabolism of lipids, carbohydrates, amino acids, cofactors, and nucleotides (**Figure 3E**). Since the anti-proliferative effects of LJH685 are negligible in A-375 cells, these results suggest that RSK inhibition reorganizes the metabolic landscape to bolster proliferation. GSEA analyses further revealed a reduction in gene sets related to hypoxia and inflammation (**Figure 3F** and **S3A**). Strikingly, the visualization of enriched GO:BP gene sets networks revealed that LJH865 led to global repression of immune response pathways except for the induction of pathways associated with antigen processing and presentation (**Figure 3G**). Detailed volcano-plot representation of these genes shows a significant induction of several MHC class I and class II transcripts in A-375 cells treated with LJH865 (**Figure 3H**). Other RSK-selective inhibitors, such as BIX02565 and LJI308, were also able to induce antigen presentation genes such as *HLA-DQA2*, *CD74*, and *HLA-DOA* (**Figure 3I-J** and **S3B**). Importantly, transcripts associated with antigen presentation were also induced in MeWo cells treated with BI-D1870 (**Figure S2A**), reinforcing a RSK-specific role. We next asked whether selective RSK inhibition could sensitize cells to IFN-*γ*, a well-established inducer of antigen presentation genes in melanoma ^35^. The induction of antigen presentation transcripts was observed in A-375 cells treated with LJH685 or IFN-*γ*, and the combination of both treatments resulted in a stronger induction (**Figure 3K**). This suggests that RSK activity status can likely modulate basal and IFN-*γ*-induced antigen presentation in melanoma tumors. As done for the study of BI-D1870 in MeWo cells (**Figure 2J**), we examined the impact of LJH685 on the expression level of transcript signatures associated with seven differentiation stages in melanoma. This RNA-Seq analysis shows that RSK markedly lowers markers of undifferentiated states while increasing neural crest-like and transitory markers (including *DCT*) (**Figure 3L**). Taken together, these results do not support the anti-proliferative and pro-inflammatory effects of specific RSK inhibition. Conversely, RSK inhibition appears to reorganize the immunometabolic landscape of melanoma cells by reducing inflammation and rewiring metabolic pathways.

**Figure 3.**
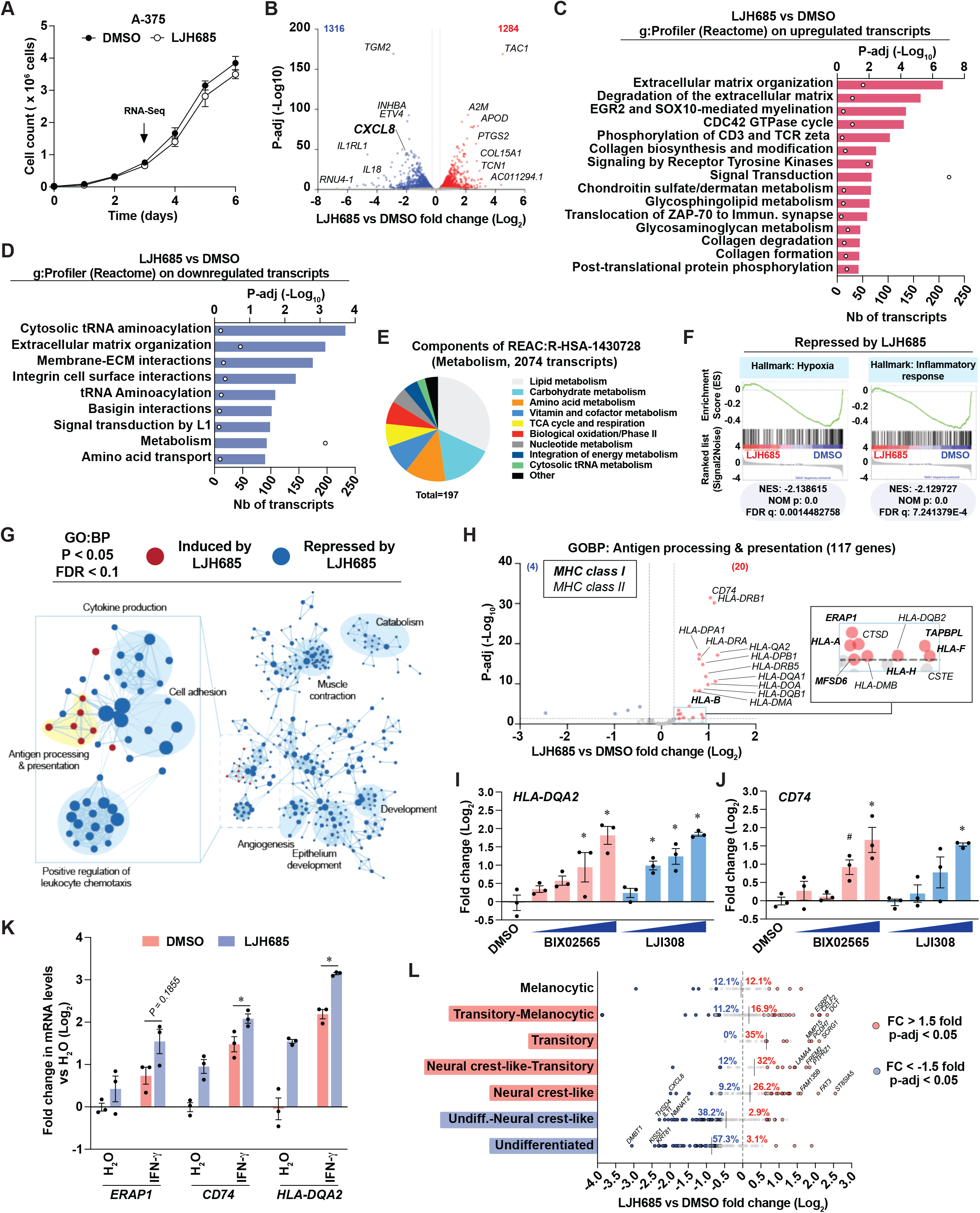
The selective RSK inhibitor LJH685 induces an anti-inflammatory gene expression program associated with enriched antigen presentation pathway transcripts. See also Figure S3. (A) Proliferation curve in A-375 cells treated with DMSO 0.1 % (v/v) or LJH685 (3 µM) for 72 h. (B) Volcano plot that indicates differentially expressed transcripts in A-375 cells treated as in A. Numbers in the upper corners indicate the number of up-regulated (red) and down-regulated (blue) significant transcripts based on threshold values of fold change > 1.2 and p-adj < 0.05. (C) Functional gene analysis (g:Profiler; Reactome) of significantly up-regulated transcripts in B. (D) Functional gene analysis as in C for significantly down-regulated transcripts in B. (E) Pie chart associating a functional category (g:Profiler) to the 197 down-regulated transcripts in the metabolism gene from D. (F) GSEA on selected significantly enriched gene sets (Hallmarks, MSigDB) on normalized data linked to panel B. NES: normalized enrichment score; NOM p: nominal p-value; FDR: False-discovery rate q-value. (G) Visualization of significantly enriched gene sets linked to panel B with Cytoscape (GO:BP, MSigDB, with p < 0.05 and FDR < 0.1). (H) Volcano plot of the antigen processing and presentation gene set (Hallmarks, MSigDB), as indicated in B. (I, J) *HLA-DQA2* and *CD74* transcript levels in A-375 cells treated with DMSO 0.1 % (v/v), BIX02565 or LJI308 (both at 0.3, 1, 3, 10 µM) for 72 h. (K) Antigen processing and presentation gene expression in A-375 cells treated with DMSO 0.1% (v/v) or LJH685 (10 µM) for 72 h and then treated with human recombinant IFN-*γ* (1 ng/mL) or water for 24 h. (L) Modulation by LJH685 in A-375 cells of transcripts associated with phenotype switching in melanoma, done on RNA-Seq data from B. The percentages indicate the proportion of significantly up-regulated or down-regulated transcripts in each gene set associated with a specific differentiation stage. All data are shown as means ± SEM from 3 independent experiments. I,J: * p < 0.05; # p < 0.1, One-Way ANOVA with Dunnett multiple comparison test (versus DMSO control). K: * p < 0.05, paired Student’s *t*-test between IFN-*γ*treatments for a given transcript.

### The pro-inflammatory effects of PLK1 targeting are linked to therapeutic resistance and phenotype switching in a cell line-dependent manner

Unselective RSK inhibitors such as BI-D1870 were shown to bind the ATP-binding site of several kinases, such as PLK1^13^. As suggested in **Figure 1A-1B** and **S1A**, targeting the expression of these kinases could impact the proliferation and pro-inflammatory response of melanoma cells and tumors. Thus, we hypothesized that the pro-inflammatory effects of BI-D1870 (**Figure 1D-1F** and **2G**) and likely BRD7389 (**Figure 1G**) are not due to RSK inhibition but likely through another kinase (**Figure S1B**). To confirm this hypothesis, we examined the induction of *CXCL8* in MeWo and A-375 cells treated for 72 h with inhibitors targeting PLK1/3, CDK2, FAK, ULK1/2, AURORA/B, CDK16, MET, and JAK3, all shown to bind BI-D1870 ^13^. While inhibition of ULK1/2 with MRT68921 had similar pro-inflammatory effects in both cell lines, all other inhibitors had distinct impacts (**Figure 4A**), as seen for BI-D1870 and BRD7389 (**Figure 1D and 1G**). The selective PLK1/3 inhibitor BI 6727 (volasertib) ^36^ was the most potent inducer of *CXCL8* in MeWo cells, with half-maximal induction with doses as low as 10 nM. In contrast to the weak effects of the RSK-selective inhibitor LJH685 on A-375 proliferation and survival (**Figure 3A** and **S1D**), BI 6727 induced cell death in A-375 with an IC_50_ of 14.1 nM (**Figure 4B**). Although MeWo cells have a similar IC_50_ (25.4 nM), a survival plateau is observed at 30-50 % relative viable cell count, as seen with Presto Blue HS and crystal violet assays (**Figure 4B-4D**). In response to BI 6727, PARP cleavage and *β*-actin degradation was higher in A-375 cells compared to MeWo cells (**Figure 4E-4G**), which suggests that cell death in A-375 cells is related to the induction of apoptosis. As an alternative approach, we performed PLK1 gene silencing in MeWo and A-375 cells (**Figure 4H**). As observed for PLK1 inhibition with BI 6727, PLK1 knockdown specifically induced *CXCL8* in MeWo cells but had no effect in A-375 cells (**Figure 4I**). In addition, PLK1 KD in A-375 induced PARP cleavage and *β*-actin degradation, supporting pro-apoptotic effects (**Figure 4J-4K**).

**Figure 4.**
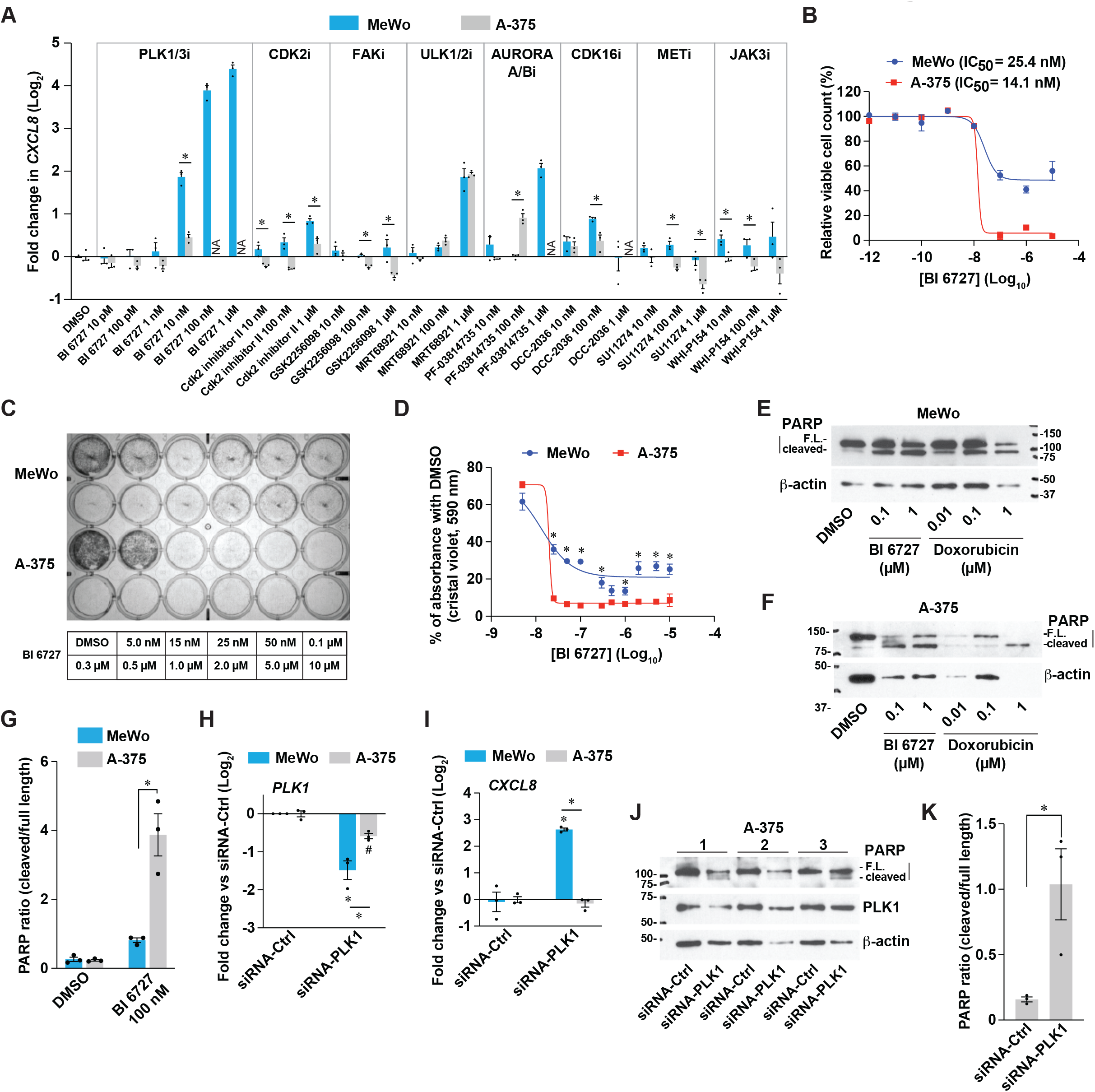
The pro-inflammatory effects of PLK1 targeting are linked to therapeutic resistance in a cell line-dependent manner. (A) *CXCL8* transcript levels in MeWo and A-375 cells treated for 72 h with DMSO 0.1 % (v/v) or a panel of inhibitors targeting putative BI-D1870 targets. NA: non-available due to cell death. (B) Viable cell counts of melanoma cells with Presto Blue HS. (C) Representative crystal violet staining of cells treated as in A. (D) Quantification of crystal violet staining, as in C. (E,F) Immunoblotting of cell extracts from cells treated as indicated for 72 h. (G) Densitometric analysis of immunoblotting from 3 independent experiments related to E and F. (H,I) *PLK1* and *CXCL8* transcript levels in cells 72 h after PLK1 knockdown. (J) Immunoblotting of cell extracts from cells treated as in H-I. Numbers indicate independent experiments. (K) Densitometric analysis of immunoblotting from 3 independent experiments related to J. All data are shown as means ± SEM from 3 independent experiments. A: * p < 0.05, unpaired Student’s *t*-test (cell lines tested separately). D,K: * p < 0.05, paired Student’s *t*-test. G-I: * p < 0.05, Two-Way ANOVA with Tukey multiple comparison test.

Next, we asked whether PLK1 targeting could modulate the expression of differentiation markers, as observed for the treatment with BI-D1870 (**Figure 2J-2L**). In MeWo cells, PLK1 inhibition repressed the levels of differentiation markers such as *TYR*, *TYRP1*, and *DCT* (**Figure 5A-5C**). The expression of the DCT protein was also fully suppressed by treatment with BI 6727 (**Figure 5D**), and PLK1 gene silencing recapitulated these effects (**Figure 5E**). To further characterize the impact of PLK1 inhibition on phenotype switching, we used the RNA-Seq data from MeWo cells treated with BI-D1870 (**Figure 2**) and identified transcripts associated with the proliferative/melanocytic and invasive/dedifferentiated states (**Figure 5F**). Testing these transcripts, we show that selective PLK1 inhibition with BI 6727 in MeWo cells induces a switch towards a more invasive/dedifferentiated state (**Figure 5G**), which is also observed with the non-selective RSK inhibitor BRD7389 (**Figure 5H**). These experiments confirm our hypothesis that non-selective RSK inhibition induces pro-inflammatory effects and dedifferentiation by inhibiting other kinases such as PLK1. These results also suggest that the cell line-specific capacity to mount a pro-inflammatory response to PLK1 inhibition is intimately linked to drug resistance and/or protection from apoptosis.

**Figure 5.**
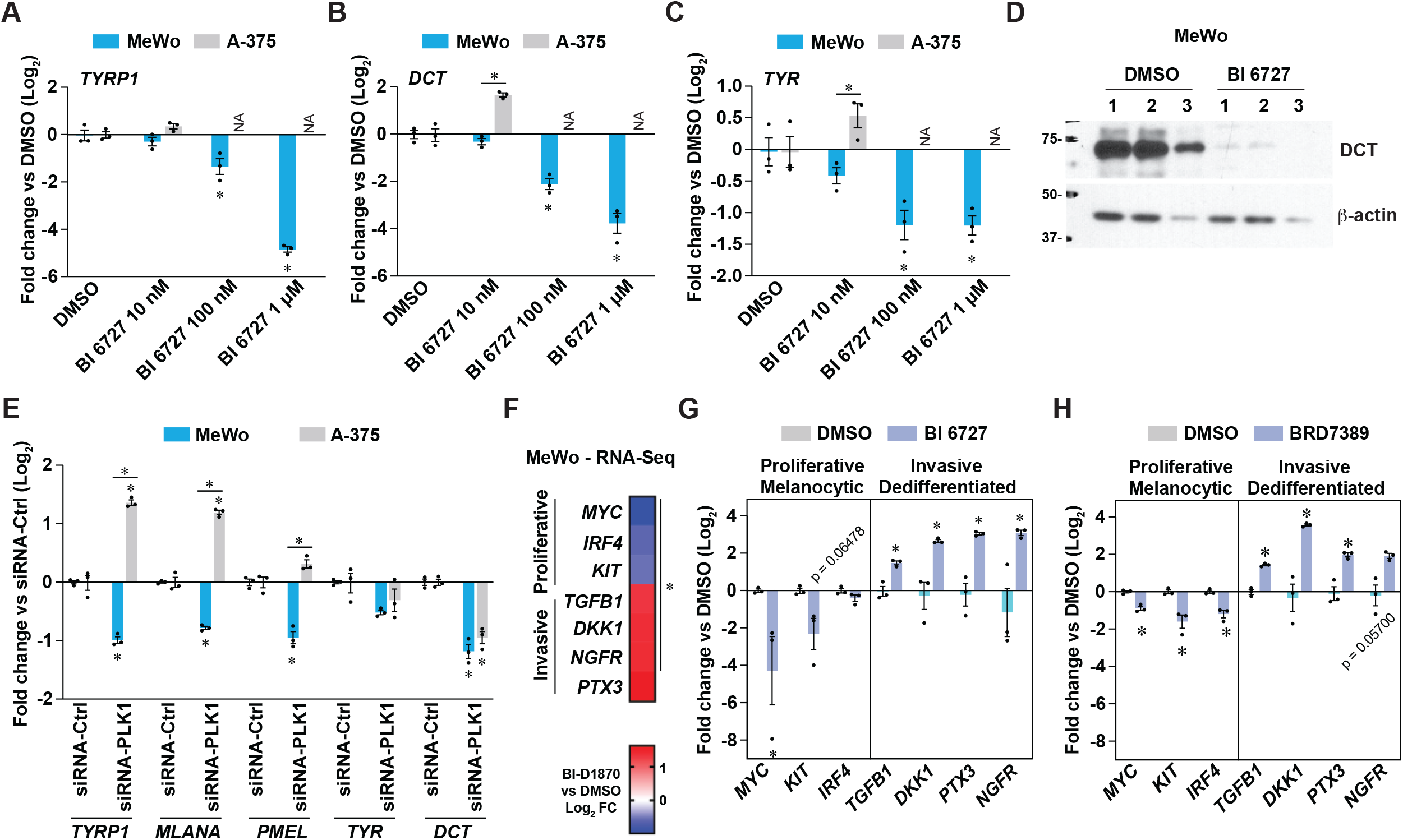
Resistance to PLK1 inhibition is associated with phenotype switching towards a dedifferentiated state in melanoma cells. (A, B, C) Gene expression in MeWo and A-375 cells treated for 72 h as indicated. NA: Not available due to cell death. (D) Immunoblotting of protein extracts from MeWo cells treated as in A-C. Numbers indicate independent experiments. (E) Gene expression of differentiation markers in cells treated with siRNA against PLK1 versus non-targeting siRNA for 72 h. (F) Heat-map of Log2 fold change of specific transcripts linked to phenotype switching in MeWo cells treated with BI-D1870 (3 µM) for 72 h. Derived from RNA-Seq analyses in Figure 2. (G,H) Gene expression in MeWo cells treated with BI 6727 (1 µM) or BRD7389 (10 µM) for 72 h. All data are shown as mean ± SEM from 3 independent experiments. A-C, E: * p < 0.05, Two-Way ANOVA with Tukey multiple comparison test. F: * p-adj < 0.05. G-H: * < 0.05, paired Student’s *t*-test.

### Mitochondrial determinants shape the sensitivity and inflammatory response to RSK and PLK1 inhibition

To determine whether a transcriptional signature is associated with the capacity of cells to resist PLK1 inhibition and mount a pro-inflammatory response, we selected melanoma cell lines based on their *CXCL8* induction pattern in response to BI-D1870 treatment (**Figure 1D**) and performed GSEA on the corresponding publicly available RNA-Seq dataset for each cell line (**Figure 6A**). These analyses revealed that pro-inflammatory cell lines (MeWo, IGR-1, SK-MEL-5) had significant enrichment in gene sets related to oxidative phosphorylation (mitochondrial respiration), lipid metabolism, and mTORC1 signaling (**Figure 6A-6B**). On the opposite, anti-inflammatory cell lines (A-375, COLO 829, Malme-3M, WM983B) showed enrichment in gene sets such as EMT, inflammation and apoptosis (**Figure 6A and 6C**). These analyses led us to hypothesize that the response to RSK and PLK1 inhibitors is defined by mitochondria, which is in line with the central role of these organelles in tumor progression and therapeutic resistance in melanoma (reviewed in ^37,38^). However, we did not find any difference that could match the two groups of cells for mitochondrial DNA (mtDNA) content (**Figure 6D**) and expression of labile respiratory complex subunits (**Figure S4A**), although some cell-specific differences were detected. Bioenergetic measurements were similar between the two groups but indicated that the A-375 cell line had distinct basal and stress-induced features compared to other cell lines. Strikingly, A-375 had a higher ratio between oxygen consumption and extracellular acidification rates (**Figure 6E**) and a higher fraction of mitochondrial respiration dedicated to ATP synthesis (**Figure 6F**). These results indicate that this cell line efficiently utilizes its mitochondria under basal conditions, which does not support basal respiration as a modulator of the response to RSK or PLK1 inhibition. However, A-375 cells were less efficient at increasing mitochondrial respiration in response to chemical uncoupling (**Figure 6G and 6H**) and were more efficient at increasing glycolysis in response to monensin, a Na^+^ ionophore that maximizes ATP hydrolysis by the Na^+^/K^+^-ATPase (**Figure 6I and 6J**). These results suggest that A-375 cells have constitutive mitochondrial configurations that make these organelles maladapted to cellular stress, which could be linked to their propensity to engage apoptosis compared to other cells. To explore the potential role of mitochondria in shaping the pro-inflammatory and resistant phenotype of MeWo cells, we used the Rho-0 protocol to generate mitochondria-incompetent cells and tested their response to PLK1 inhibition. Compared with parental cells, Rho-0 cells were characterized by dramatic reduction in the mitochondria DNA/genomic DNA ratio, reduced expression of labile respiratory proteins subunits, and the absence of mitochondrial respiration (**Figure 6K**, **S4B and S4D**). In addition, Rho-0 cells showed a compensatory increase in glycolysis (**Figure 6K** and **S4C**). Strikingly, Rho-0 cells showed major changes in their sensitivity profile to BI 6727 treatment compared to MeWo cells (**Figure 6L**). While being less apoptotic in response to BI 6727 at lower doses (**Figure S4E and S4F**), Rho-0 cells lost the resistance to apoptosis at 10 µM BI 6727 (**Figure 6L and 6M**), a dose from the survival plateau in MeWo cells (**Figure 4B and 4D**). Rho-0 cells were also less potent at inducing *CXCL8* and *IL6* in response to RSK and PLK1 inhibitors (**Figure 6N and 6O**). Taken together, these results suggest that specific expression patterns of mitochondrial proteins may be responsible for the sensitivity and inflammatory profiles of melanoma cells in response to RSK and PLK1 inhibitors.

**Figure 6.**
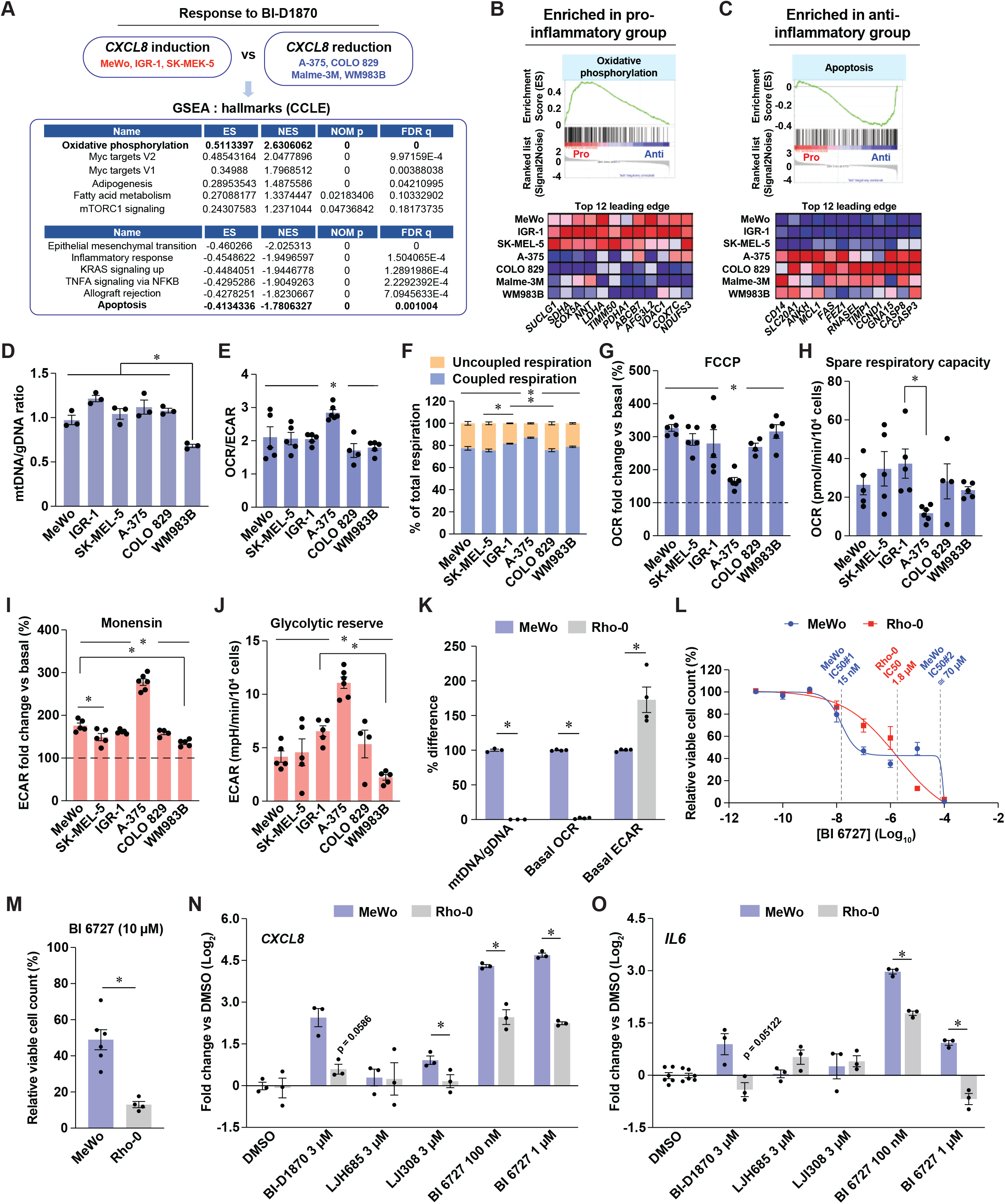
Mitochondrial determinants shape the sensitivity and inflammatory response to RSK and PLK1 inhibition. See also Figure S4. (A) Top gene set enrichments identified by GSEA in transcriptomic datasets (CCLE, Broad 2019) of human melanoma cells subsets defined by their response to BI-D1870. ES: enrichment score; NES: normalized enrichment score; NOM p: nominal p-value; FDR q: false discovery rate q-value. (B, C) Details for gene set enrichment analyses and top 12 leading edge transcripts of selected gene sets from A. Pro/Anti: cell group with pro-or anti-inflammatory response to BI-D1870. (D) Mitochondrial DNA to genomic DNA ratio in human melanoma cell lines. (E) Oxygen consumption rate (OCR) to extracellular acidification rate (ECAR) ratio in human melanoma cell lines. (F) Metabolic organization in melanoma cell lines. (G) Maximal induction of OCR by acute FCCP treatment relative to basal conditions in human melanoma cell lines. (H) Spare respiratory capacity in human melanoma cell lines. (I) Maximal induction of ECAR by oligomycin and monensin relative to basal conditions in melanoma cells. (J) Glycolytic reserve in human melanoma cell lines. (K) Characterization of Rho-0 MeWo cells by comparison of mitochondrial features with parental MeWo cells. (L) Relative viable cell counts with Presto Blue HS. Cells were treated for 72 h with 10-fold dilutions of BI 6727. (M) Details of viable cell count at 10 µM BI 6727, related to L. (N,O) *CXCL8* and *IL6* transcript levels in MeWo and MeWo-Rho-0 cells treated DMSO 0.1 % (v/v) and inhibitors at the indicated doses for 72 h. All data are shown as means ± SEM from 3 or more independent experiments. D-J: * p < 0.05, One-Way ANOVA with Tukey multiple comparison test; K, M-O: * p < 0.05, paired Student’s *t*-test, except for mtDNA/gDNA data in K (unpaired Student’s *t*-test).

### Defining actionable mitochondrial signatures associated with resistance to PLK1 targeting in melanoma

To characterize mitochondrial protein expression patterns associated with resistance to PLK1 inhibition in melanoma cell lines, we performed correlation analyses between transcript expression levels (CCLE) and the impact of PLK1-targeting approaches on cell proliferation (DepMap). PLK1 targeting approaches included gene knockout by CRISPR/Cas9, gene silencing by RNA interference, and PLK1 inhibitors: BI 6727, BI 2536, GSK461364, HMN-214, NMS-1286937, and GW-843682X (**Figure 7A**). For the selection of correlating transcripts with good confidence, we chose transcripts that showed a significant correlation with the proliferation impact for at least 2 PLK1-targeting approaches. This led to the primary identification of 881 and 1501 transcripts that respectively correlate positively and negatively with resistance to PLK1-targeting (**Figure 7B**). Gene functional classification of ‘’sensitizing’’ transcripts (negative correlation coefficient *ρ*) revealed enrichment for translation, ribosome biogenesis, and cell division pathways, while ‘’resistance’’ transcripts (positive *ρ*) were enriched for development, metabolic, and pigmentation pathways (**Figure 7C**). Considering that the A-375 cell line is more proliferative than the MeWo cell line (**Figure S5A**), these analyses show that the sensitivity to PLK1 targeting is higher in cell lines more committed to cell division, which is expected given the role of PLK1 in this process. However, these analyses also indicate that metabolic pathways are associated with resistance to PLK1 targeting. In line with our hypothesis that mitochondrial determinants are linked to this resistance, we next identified 124 transcripts encoding mitochondrial proteins that significantly correlate with the sensitivity to PLK1 targeting in melanoma cells. By matching each gene with the gene essentiality (CRISPR/Cas9) data from the DepMap portal, we found that a major fraction of ‘’sensitizing’’ and ‘’resistance’’ genes are classified as nonessential (**Figure 7D**), which opens the possibility of exploiting the targeting of these genes for experimental and therapeutic purposes. All six PLK1 inhibitors had a highly similar correlating pattern with most genes from the mitochondrial signature (**Figure 7E**). Interestingly, the unselective RSK inhibitor BI-D1870 clustered with both PLK1 knockdown and PLK1 inhibitors, further supporting the inhibition of PLK1 by this compound. Functional classification of the gene signature as listed by MitoPathways3.0 revealed that sensitizing transcripts were associated with mitochondrial central dogma (replication, transcription, translation), carbohydrate metabolism and apoptosis, while resistance transcripts were associated with lipid metabolism, amino acid metabolism and detoxification (**Figure 7F**). Using STRING, we also performed a protein-protein interaction analysis of the whole mitochondrial transcript signature. This analysis shows that most proteins encoded by the signature are part of a tight interaction network that parallels functional classification analyses such as mitochondrial central dogma, mitochondrial metabolism, OXPHOS, and apoptosis (**Figure 7G**). To test whether the targeting of resistance transcripts can indeed affect the response to PLK1 inhibition, we silenced three of these transcripts in MeWo cells, *ABCD1* (ATP-binding cassette sub-family member 1), *BCL2A1* (Bcl-2-related protein A1), and *PRKACA* (cAMP-dependent protein kinase catalytic subunit *α*), chosen for their strong correlation with inflammatory transcript clusters in human melanoma tumors (**Figure S5B-S5F**). Individual knockdown of these three transcripts reduced the induction of *CXCL8* and *IL6* by BI 6727 treatment, with siRNA targeting *ABCD1* being the most potent treatment (**Figure 7H**). *ABCD1* knockdown also lowered the resistance plateau previously observed in MeWo cells treated with high doses of BI 6727 (**Figure 7I**). Taken together, these analyses show that mitochondrial signatures are associated with distinct responses to RSK and PLK1 inhibitors (**Figure 7J**), and that targeting specific mitochondrial proteins is a valid approach to fine-tune the sensitivity of melanoma cells to PLK1 inhibition.

**Figure 7.**
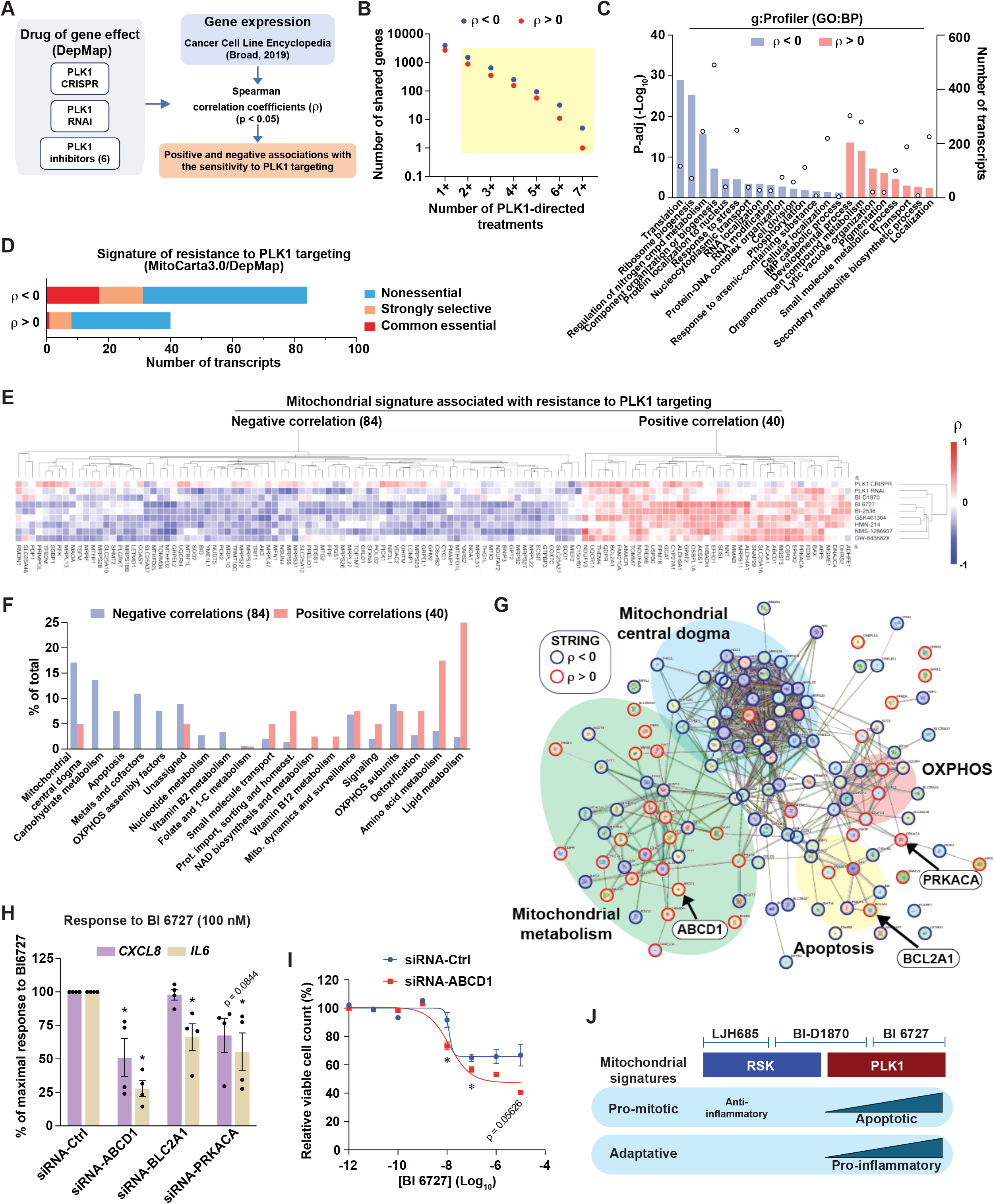
Defining actionable mitochondrial signatures associated with resistance to PLK1 targeting in melanoma. See also Figure S5. (A) Schematic depicting the strategy to identify transcripts associated with resistance to PLK1 targeting. The six PLK inhibitors, in addition to BI-D1870, are indicated in panel E. (B) Transcripts that significantly correlate (p < 0.05) with 2 or more PLK1-directed treatments were selected for further analyses. (C) Functional gene analysis (g:Profiler, GO:BP) performed on correlating transcripts from B. Bars indicate p-adj values and circles indicate number of transcripts associated with a gene set. (D) Essentiality of transcripts encoding for mitochondrial proteins that significantly correlate with resistance to PLK targeting in melanoma cells (DepMap). (E) Heat-map representation and hierarchical clustering of Spearman correlations between the 124 mitochondrial signature transcripts and respective impact on proliferation dynamics of BI-D1870 and PLK1 targeting approaches (DepMap). (F) Functional classification of 124 mitochondrial signature transcripts associated with resistance to PLK1 targeting, according to MitoPathways3.0. (G) STRING interaction network of the 124 mitochondrial signature transcripts associated with resistance to PLK1 targeting. Nodes indicate proteins, and edges indicate interactions. Spearman’s correlation with proliferation dynamics (DepMap) is indicated by nodes’ colored borders. (H) Gene expression in MeWo cells transfected with the indicated siRNA for 48 h and treated with DMSO 0.1 % (v/v) or BI 6726 (100 nM) for 72 h. (I) Relative viable cell counts of MeWo cells treated as in H but with 10-fold dilutions of BI 6727, using Presto Blue HS. (J) Model depicting the impact of mitochondrial signatures on the response to RSK and PLK inhibitors. All data are shown as means ± SEM from 3 or more independent experiments. * p < 0.05, One-Way ANOVA with Dunnett multiple comparison test.

## DISCUSSION

The first evidence of PLK1 as a putative target for melanoma treatment emerged from seminal studies showing increased PLK1 expression in melanoma compared to normal tissue and melanocytes, and lower survival of patients with high tumoral expression, also suggesting its prognostic value ^19,20^. The specific induction of apoptosis in melanoma cells versus melanocytes ^39^ has reinforced the interest in targeting PLK1 with small molecules (BI 2536 ^40,41^, BI 6727 ^42^) and launching clinical trials (BI 6727 ^43^ and other inhibitors, reviewed in ^21^). Over the past years, PLK1 inhibition appeared as an attractive option for melanoma treatment in combination with MEK inhibition ^44–46^, NOTCH inhibition ^47^, or in the case of resistance to directed therapies (BRAF and MEK inhibition ^48^). PLK1 inhibitors appear as versatile anticancer agents for a variety of cancers, and their recent interplay with immunotherapies ^49,50^ will likely open interesting research avenues for melanoma. While PLK1 inhibition in experimental models shows potent anti-tumoral effects in vitro and in vivo, the translation to the clinic has been curbed by modest anti-tumor activity, tumor relapse, and the frequent lack of response to PLK1 inhibitors ^51–55^. Very few studies examined the factors that could confer resistance to PLK1 inhibition. Mutations in the ATP-binding site of the PLK1 gene ^56,57^, increased expression of the AXL/TWIST1/MDR1 ^56,58^, and the induction of mitotic slippage at high doses of PLK1 inhibitors ^59^ are among resistance mechanisms observed after the establishment of resistant cell lines. Herein, we show that some melanoma cell lines, such as MeWo, possess intrinsic resistance toward PLK1 inhibition without the need to establish resistant cell lines. In response to PLK1 inhibition, MeWo cells become non-proliferative and mount a potent pro-inflammatory response. From a clinical perspective, these observations suggest that PLK1 inhibition may quickly lead to resistant satellites associated with immunological remodeling of the tumor microenvironment. To our knowledge, our study is the first to highlight the role of mitochondria in establishing cell-intrinsic resistance to PLK1 inhibition.

The role of mitochondria in cancer has long been overlooked due to the propensity of cancer cells to divert cellular bioenergetics towards aerobic glycolysis (Warburg effect). This simplistic view of mitochondria shaded the multiple roles of this organelle in many cellular processes beyond oxygen consumption and ATP synthesis (reviewed in ^60^). Importantly, mitochondria can potently shape the response to therapies through a variety of mechanisms ^37,61,62^. Here, we found that mitochondria shape the response of melanoma cells to PLK1 inhibition by acting like a switch between immunologically silent cell death and pro-inflammatory survival. We have identified subsets of nuclear genes that encode mitochondrial proteins that can either enhance or decrease the impact of PLK1-directed treatments (pharmacological inhibition, gene silencing, gene knockout) on melanoma cell proliferation. Recently, a genome-wide CRISPR screen allowed the identification of genes that modulate the sensitivity to PLK1 inhibition, among which genes involved in chromosome attachment and segregation on the mitotic spindle were identified as sensitizers ^63^. We examined the data from this study to determine whether our candidate genes for a mitochondrial resistance signature were included ^63^. Interestingly, the knockout of the ‘’resistance’’ gene *PRKACA* sensitized cells to PLK1 inhibition, while the knockout of 15 ‘’sensitivity’’ genes from our signature rescued cell proliferation. Although this study used a non-melanoma cell line, the NALM-6 pre-B lymphocytic leukemia suspension cell line, these findings may suggest that mitochondria are general modulators of the response to PLK1 inhibition. *PRKACA* encodes for the catalytic subunit *α*of PKA, a protein kinase that has pleiotropic roles in metabolism, inflammation, and apoptosis. In melanoma, PKA can modulate BRAF and CRAF ^64,65^, and overexpression of PRKACA induces resistance to MAPK inhibition ^66^. PKA has several substrates on the outer mitochondrial membrane and within mitochondria and thus can modulate several processes such as OXPHOS, apoptosis, mitochondrial dynamics and mitophagy (review in ^67^). Although PKA appears as a candidate target for melanoma treatment, no selective inhibitor against this kinase is currently available. Among the other ‘’resistance’’ genes we have identified, BCL2A1/Bfl-1 encodes for a Bcl-2-related anti-apoptotic protein that interacts with multiple pro-apoptotic proteins and blocks the release of cytochrome c (reviewed in ^68^). In melanoma, it has been shown that this gene is regulated transcriptionally by MITF and that its expression is associated with resistance to BRAF inhibition ^69^.

While no highly selective BCL2A1 inhibitors have been identified yet ^70^, a few inhibitors have shown promising results in experimental cancer studies ^69,71^, including the BET inhibitor CPI-0610, which indirectly targets the transcription of *BCL2A1* ^72^. Our analyses also revealed that the ABC transport ABCD1 is linked to resistance to PLK1 targeting in melanoma. ABCD1 is critical for the peroxisomal import of very long fatty acids and related CoA esters for catabolism through *β*-oxidation, and mutations of this gene are responsible for adrenoleukodystrophy, an X-linked and neurodegenerative disorder associated with microglial apoptosis ^73^. In astrocytes and oligodendrocytes, the silencing of ABCD1 impaired mitochondrial respiration and induced mitochondrial superoxide anion production^74^, which suggests that this gene could exert protective and anti-apoptotic functions in melanoma cells. Aside from these precision oncology approaches, our study also shows that the depletion of mtDNA in MeWo cells dramatically alters the resistance profile of MeWo cells to PLK1 inhibition. Rho-0 cells still possess mitochondria, but they are part of fragmented networks and show no respiratory activity. Since our data indicate that OXPHOS is not associated with resistance, it would be interesting to determine which of the genes from our PLK1-targeting resistance signature are expressed in Rho-0 cells and if this model can be used to generate resistant cells. Interestingly, Rho-0 cells were more resistant than parental MeWo cells to low doses of PLK1 inhibitor. We believe that this resistance may be linked to the slower division rate of Rho-0 cells, which is in line with PLK1 inhibition being more potent in rapidly dividing cells.

In this study, we also show that the anti-proliferative and pro-inflammatory effects of some RSK inhibitors are likely due to the inhibition of other kinases, such as PLK1. Strikingly, BI-D1870 and BI 6727 have similar pro-inflammatory effects in MeWo cells, but they are respectively anti-inflammatory and pro-apoptotic in A-375 (**Figure 1D and 4A**). This suggests that the anti-inflammatory effects of BI-D1870 in A-375 (**Figure 1D**) are linked to RSK inhibition, as observed with selective RSK inhibitors such as LJH685 and LJI308 (**Figure 1D and 1H**). Indeed, despite its impact on PLK1 or other kinases, treatment with BI-D1870 also shares with LJH685 the induction of antigen presentation genes (**Figure S2A and 3G-K**). One important observation of our study is that specific inhibition of RSK does not typically induce *CXCL8* and has rather mild effects on proliferation. We have previously shown that RSK inhibitors can reduce glycolytic flux through direct phosphorylation of PFK-2 ^75^. Since RSK is involved in the control of mRNA translation and is also a potent modulator of inflammation through the TRAF6-IKK-NF-*κ*B pathway ^76–78^, we hypothesize that RSK inhibition in melanoma may globally reduce the bioenergetic charge and associated oxidative stress in melanoma cells. Interestingly, RSK targeting by RNA interference was shown to suppress proliferation in vitro and in vivo ^22,75^. While context-dependent roles of RSK may explain these discrepancies, kinase-independent functions of RSK through inhibitory interactions with CBP and ERK may explain differences between RSK inhibition and gene/RNA silencing^79,80^.

Tumor heterogeneity constitutes a major obstacle to the success of targeted therapies for the treatment of melanoma. By associating mitochondrial protein expression levels with the ability to resist PLK1 inhibition, our study reveals the existence of mitochondrial signatures participating in tumor heterogeneity. Characterizing these mitochondrial signatures associated with resistance will make it possible to predict the effectiveness of targeted therapies and design new mitochondrial therapies.

## ACKNOWLEDGMENTS

This work was supported by the Cancer Research Society [Grant 840633, 2021, Grant 1054571, 2023] and the National Sciences and Engineering Research Council of Canada [Grant RGPIN-2023-03828, 2023]. The authors thank the IRIC genomic platform and IRIC bioinformaticians for assistance with RNA-Seq. The authors thank the Centre for Applied Genomics (The Hospital for Sick Children, Toronto) for STR profiling. The authors thank Caroline Thivierge, Hamza Haddouch and Rita Maria Kenaan El Rahbani for editing the manuscript.

## AUTHOR CONTRIBUTIONS

**Émilie Lavallée:** Methodology, Investigation, Validation, Writing – Original Draft. **Maëline Roulet-Matton:** Investigation, Validation. **Viviane Giang:** Investigation, validation. **Roxana Cardona Hurtado:** Investigation, validation. **Dominic Chaput:** Investigation, validation. **Simon-Pierre Gravel:** Conceptualization, Methodology, Investigation, Formal analysis, Visualization, Writing – Original Draft, Writing – Review & Editing, Supervision, Project administration, Funding acquisition.

## MATERIALS AND METHODS

### Lead contact and material availability

Further information and requests should be directed to the Lead Contact, Simon-Pierre Gravel.

### Cell lines and treatments

Human melanoma cell lines A-375, MeWo, SK-MEL-5, IGR-1, RPMI-7951, COLO 829, Malme-3M, WM983B, and Mel JuSo were kindly provided by Pr. Ian Watson (McGill University). All cell lines, including MeWo-Rho-0, were authenticated by comparing STR profiling with the ATCC and Cellosaurus databases. All cell lines tested negative for mycoplasma using a PCR mycoplasma detection kit (#G238, Applied Biological Materials Inc., Richmond, BC, Canada). Cell lines were cultured in RPMI-1640 medium (#350-000-CL, Wisent Inc., St-Jean-Baptiste, QC, CAN) supplemented with 10 % fetal bovine serum (FBS, #090-150, Wisent Inc.) and 1 % Penicillin/Streptomycin (#450-201-EL, Wisent Inc.). Cells were maintained at 37 °C in a humidified incubator with 5 % CO_2_. Cells were tested negative for mycoplasma using a PCR detection kit (#G238, Applied Biological Materials Inc.). Rho-0 cells were generated from parental MeWo cells cultured in media containing 50 µg/mL uridine (#URD222, Bioshop Canada Inc., Burlington, ON, CAN), 100 µg/mL sodium pyruvate (#600-110-EL, Wisent Inc.), and 100 ng/mL ethidium bromide (#ETB333, Bioshop Canada Inc.) for several weeks. Rho-0 media was replaced with regular media a few days before experiments.

Inhibitors and compounds were resuspended in DMSO (#BP231-100, Fisher Scientific, Waltham, MA, USA), as recommended by the manufacturer, and stored in a -80 °C freezer. Before experiments, compound stocks were further diluted in DMSO to obtain a final DMSO concentration of 0.1 % (v/v) in culture media. BI-D1870 (#15264), BI 6727 (#18193), BIX 02565 (#19183), BRD7389 (#20214), Cdk2 inhibitor II (#15154), DCC-2036 (#21465), GSK2256098 (#22995), LJH685 (#19913), LJI308 (#19924), MRT68921 (#19905), PF-03814735 (#15015), SU11274 (#14861), and WHI-P154 (#16178) were purchased from Cayman Chemical (Ann Arbor, MI, USA). Doxorubicin hydrochloride (ab120629) was purchased from Abcam Inc. (Toronto, ON, CAN). Recombinant human interferon-gamma (IFN-*γ*, #300-02) was from PeproTech (Cranbury, NJ, USA) and stored at 100 µg/mL (10 % (v/v) fetal bovine serum in sterile water) in a -80 °C freezer.

### Gene silencing

A-375 and MeWo cells were seeded on 6-well culture plates with 2 mL of media per well and at cell densities of 15,000 cells/mL and 75,000 cells/mL, respectively. Cells were incubated for 16-24 h before transfection with siRNA. Non-targeting siRNA (Allstar, 1027280) and siRNA pools, made of four siRNAs (FlexiTube Gene Solutions, 1027416) against human PLK1 (GS5347), ABCD1 (GS215), BCL2A1 (GS597), and PRKACA (GS5566) were purchased from Qiagen (Hilden, DEU). Cells were transfected with Lipofectamine RNAiMAX transfection reagent (#13778500, ThermoFisher Scientific, Waltham, MA, USA) using Opti-MEM low serum medium (#31985070, ThermoFisher Scientific). The final concentration of the non-targeting control siRNA and siRNA pools was 10 nM (2.5 nM for each siRNA used for pools), except PLK1, which was 1 nM. Transfections were done following the manufacturer’s protocol, using 5 µL of transfection reagent per well. Media were replaced 24 h post-transfection.

### Cell counting and viability assay

Cells were seeded on 6-well culture plates in 2 mL of media at the following densities: 9,375 cells/mL (A-375), 50,000 cells/mL (Mel JuSo, SK-MEL-5), 75,000 cells/mL (MeWo, Malme-3M), 70,000 cells/mL (RPMI-7951), 62,500 cells/mL (COLO 829), 30,000 cells/mL (IGR-1) and 25,000 cells/mL (WM983B). Cells were incubated for 16-24 h before treatments. 72 h post-treatment, cells were rinsed with PBS 1X (#311-010-CL, Wisent Inc., St-Jean-Baptiste, QC, CAN), dissociated with Trypsin/EDTA (0.25 % Trypsin and 2.21 mM EDTA 4Na, #325-043-EL, Wisent Inc.) and resuspended in 1 mL of RPMI-1640 medium (#350-000-CL, Wisent Inc.). Cell suspensions were mixed with trypan blue solution (0.4 % solution in phosphate buffer saline, #609-130-EL, Wisent Inc.) and counted manually with a hemacytometer (#1492, Hausser Scientific, Horsham, PA, USA). For proliferation curves, cell culture media was changed every day with the renewal of treatments. Occasionally, cell suspensions were centrifuged at 1000 x g for 5 min and resuspended in a smaller volume of culture medium.

For the assessment of viability, cells were seeded in 96-well plates at 1,250 cells/100 µL (A-375), 7,500 cells/100 µL (MeWo), 10,000 cells/100 µL (Rho-0) to which 100 µL of 2X concentrated inhibitor dilutions (BI-D1870, LJH685, and BI 6727) were added. Six wells per plate contained only cell culture media (no cell control). Cell viability was measured 72 h later with the PrestoBlue HS cell viability reagent (#P50201, ThermoFisher Scientific, Waltham, MA, USA) following the manufacturer’s protocol. Absorbance was measured with a Varioskan LUX multimode microplate reader (ThermoFisher Scientific). For crystal violet staining, cells were seeded on 24-well culture plates in 500 µL of media at the following densities: 7,500 cells/mL (A-375), and 45,000 cells/mL (MeWo). Cells were incubated for 16–24 h before treatments. 72 h post-treatment, cells were rinsed with PBS 1X and fixed with 10% formalin (v/v in water) for 15 min. Cells were rinsed with water and stained with 0.5% crystal violet solution (w/v in 20% methanol) for 30 min. Cells were rinsed with water and plates were allowed to dry completely. Pictures were taken with the ChemiDoc XRS+ using the Image Lab software (Bio-Rad, Hercules, CA, USA). Dye extraction was done with 33% (v/v) acetic acid solution for 30 min and absorbance at 590 nm was measured with a Varioskan LUX multimode microplate reader (ThermoFisher Scientific).

### Gene expression analysis

Total cellular RNA was isolated using the Monarch Total RNA Miniprep Kit (#T2010, NEB, Ipswich, MA, USA) following the manufacturer’s protocol. RNA concentration was obtained using a Varioskan LUX multimode microplate reader (ThermoFisher Scientific, Waltham, MA, USA) with a µDrop plate (ThermoFisher Scientific). Reverse transcription was performed with the LunaScript RT SuperMix Kit (#E3010, NEB) using the manufacturer’s protocol on a SimpliAmp thermal cycler (Applied Biosystems, Waltham, MA, USA). Oligonucleotide primers were conceived with the Primer Blast software (NCBI, MD, USA), synthesized by Integrated DNA Technologies (IDT) (Coralville, IA, USA), and validated for efficiency and specificity. The list of primers for the detection of human transcripts by qRT-PCR is the following (5*′*à3*′*direction):

*TBP* forward: TGCCACGCCAGCTTCGGAGA

*TBP* reverse: ACCGCAGCAAACCGCTTGGG

*UBE2D2* forward: AGAGAATCCACAAGGAATTGAATGA

*UBE2D2* reverse: TAGGGACTGTCATTTGGCCC

*CXCL8* forward: TGATTTCTGCAGCTCTGTGT

*CXCL8* reverse: AAACTTCTCCACAACCCTCT

*IL6* forward: CAGAGCTGTGCAGATGAGTA

*IL6* reverse: GCGCAGAATGAGATGAGTTG

*HLA-DOA* forward: GGGGTTCCACACCCTGATGA

*HLA-DOA* reverse: CGCCGTAAGACTGGTAGAAGG

*HLA-DQA2* forward: CCCTGTGGAGGTGAAGACAT

*HLA-DQA2* reverse: AGTCTCTTTCGTCTCCAGGTC

*CD74* forward: TTGGAGCAAAAGCCCACTGA

*CD74* reverse: GCACTTGGGCCTGAATGAAC

*ERAP1* forward: TTTCACTTTCGGTCCTGGGG

*ERAP1* reverse: TGAGGGGCAGAAACACCATC

*PLK1* forward: AGTACGGCCTTGGGTATCAG

*PLK1* reverse: GCTCGCTCATGTAATTGCGG

*BCL2A1* forward: ACCAGGCAGAAGATGACAGA

*BLC2A1* reverse: TGGTATCTGTAGGACGCACT

*ABCD1* forward: ACTCAGTGGAGGACATGCAA

*ABCD1* reverse: ACGTCCTTCCAGTCACACAT

*PRKACA* forward: CGAGCAGGAGAGCGTGAAAGA

*PRKACA* reverse: CCAAGTGGGCTGTGTTCTGA

*TYRP1* forward: CATGCAGGAAATGTTGCAAGAA

*TYRP1* reverse: AGTTTGGGCTTATTAGAGTGGAATC

*DCT* forward: GTCTGTGGCTCTCAGCAAGG

*DCT* reverse: ATAGCCGGCAAAGTTTCCTGT

*TYR* forward: AGCTATCTACAAGATTCAGACCCAG

*TYR* reverse: TGACGACACAGCAAGCTCA

*PMEL* forward: TCCAGGCTTTGGTTCTGAGT

*PMEL* reverse: TGTGATAGGTGCTTTGCTGG

*MLANA* forward: GCTCATCGGCTGTTGGTATT

*MLANA* reverse: AGAGACACTTTGCTGTCCCG

*MYC* forward: TACTGCGACGAGGAGGAGAA

*MYC* reverse: CGAAGGGAGAAGGGTGTGAC

*IRF4* forward: CCCGGAAATCCCGTACCAAT

*IRF4* reverse: AGGTGGGGCACAAGCATAAA

*KIT* forward: AACACGCACCTGCTGAAATG

*KIT* reverse: GTCTACCACGGGCTTCTGTC

*TGFB1* forward: CGCGTGCTAATGGTGGAAAC

*TGFB1* reverse: GCTGAGGTATCGCAAGGAAT

*DKK1* forward: TCACGCTATGTGCTGCCC

*DKK1* reverse: TGACCGGAGACAAACAGAACC

*NGFR* forward: AGGGAGGAATCGAGGAACCA

*NGFR* reverse: TCTGGCTTTGGGCGAATCAT

*PTX3* forward: AGTGCCTGCATTTGGGTCAA

*PTX3* reverse: CTCTCCACCCACCACAAACA

Quantitative RT-PCR was done with the Luna Universal qPCR Master Mix (#M3003, NEB,) using a QuantStudio 5 Real-time PCR system (Applied Biosystems). PCR program: 95 °C for 3 min followed by 45 cycles at 95 °C for 15 sec and 60 °C for 30 sec. A melting curve analysis was performed, consisting of 95 °C for 15 sec, 60 °C for 1 min and an increase in temperature back to 95 °C at 0.1 °C/s pace. The QuantStudio Design and Analysis desktop software (Applied Biosystems) was used for data analysis. Gene expression level was obtained using the ΔΔCt calculation method, where RNA levels were normalized against *TBP* or *UBE2D2*.

### Quantification of mitochondrial DNA

Cells were lysed in SNET lysis buffer (20 nM Tris-HCl pH 8.0, 5 mM EDTA pH 8.0, 400 mM NaCl, 1 % (w/v) SDS, 100 µg/mL RNAse A, and 400 µg/mL proteinases K) for 16-20 h at 55 °C on a digital block heater. Cellular DNA was extracted with one volume of phenol:chloroform:isoamyl alcohol (25:24:1, #15593031, ThermoFisher Scientific, Waltham, MA, USA), precipitated with one volume of isopropyl alcohol, and centrifuged at 17,000 x g for 10 min. Pellets were washed with 70 % ethanol and allowed to dry at room temperature before being dissolved in TE buffer (1 mM EDTA pH 8.0, 10 mM Tris-HCl pH 8.0) for 16-20 h at 4 °C. The same procedure was used for sample preparation for STR profiling. DNA concentrations were measured with a Varioskan LUX multimode microplate reader (ThermoFisher Scientific). Mitochondrial and genomic DNA were quantified by quantitative PCR as indicated in Gene expression analysis. The PCR program was 95°C for 6 min followed by 45 cycles of 95 °C for 15 sec and 60 °C for 30 sec. The following list of primers was used to amplify human mitochondrial and genomic DNA (5→3*′*direction):

*MT-CYB* (mitochondrial DNA) forward: GCGTCCTTGCCCTATTACTATC

*MT-CYB* (mitochondrial DNA) reverse: CTTACTGGTTGTCCTCCGATTC

*RPL13A* (genomic DNA) forward: CTCAAGGTCGTGCGTCTG

*RPL13A* (genomic DNA) reverse: TGGCTTTCTCTTTCCTCTTCTC

### Western blotting

Cells were washed with ice-cold PBS and lysed with RIPA low SDS buffer (50 mM Tris-HCl pH 7.4, 150 mM NaCl, 5 mM EDTA pH 8.0, 1 mM EGTA pH 8.0, 1 % NP-40, 0.5 % sodium deoxycholate, 0.1% SDS) supplemented with sodium fluoride, sodium orthovanadate, and protease inhibitor cocktail (ab270055, Abcam Inc.,Toronto, ON, CAN). Lysates were frozen and thawed 3 times in liquid nitrogen to maximize lysis efficiency and were kept on ice for 30 min. Samples were centrifuged at 17,000 x g for 10 min at 4 °C. Protein concentration was determined with the Pierce BCA Protein Assay Kit (#23225, ThermoFisher Scientific, Waltham, Ma, USA). Equal amounts of total proteins (10 μg) were denatured in sample buffer (50 mM Tris-HCl pH 6.8, 100 mM DTT, 2 % SDS, <1.5 mM bromophenol blue 1.5 mM, 1.075 M glycerol) at 95 °C for 5 min. The denatured proteins were resolved on 8–12 polyacrylamide-SDS gels and transferred onto PVDF membrane (GE Healthcare, Chicago, IL, USA). A molecular weight marker was added onto each gel for reference (Precision Plus Protein Dual Color, #1610374, Bio-Rad, Hercules, CA, USA). Transfer on PVDF membranes was done using Pierce Western blot transfer buffer (#35040, ThermoFisher Scientific) and Trans-Blot SD semi-dry transfer cell (Bio-Rad, Hercules, CA, USA) at 22 V and 0.2 A for 50 min. Blots were blocked in 5 % non-fat dry milk diluted in tris buffer saline (TBS) supplemented with 0.1% Tween-20 (TBST) and incubated with primary antibodies overnight at 4 °C with gentle rocking. Primary antibodies against PARP (#9542), *β*-actin (#3700), PLK1 (#4535), and Melan-A/MART-1/MLANA (#64718) were purchased from Cell Signaling Technology Inc. (Whitby, ON, CA). Antibodies against OXPHOS protein complexes (#45-8099), Vinculin (#MA5-11690) were purchased from ThermoFisher Scientific. Antibodies against Dopachrome Tautomerase/DCT (#HPA010743) and Tyrosinase/TYR (05-647) were from MilliporeSigma (Oakville, ON, CAN). Membranes were washed 5 times for 5 min with TBST and then incubated with anti-mouse or anti-rabbit secondary antibodies coupled with horseradish peroxidase at 1:10,000 dilution (#KP-5220-0341, #KP-5220-0336, Mandel Scientific, Guelph, ON, CAN) at room temperature for 1 h. Samples were revealed using the SuperSignal West Pico PLUS Chemiluminescent substrate (#34579, ThermoFisher Scientific). Chemiluminescence was detected on Blu-Lite films (Dutscher, Bernolsheim, FR). Films were developed in a dark room using an X-ray film processor. Densitometric analyses were done using the ImageJ software (NIH).

### Cytokine array

MeWo cells were seeded in 6-well plates (2 mL/well) at a cell density of 75,000 cells/mL. Cells were incubated for 16-24 h before treatments. 48 h post-treatment, the media was replaced by new media with the same concentration of the corresponding inhibitor. Conditioned media from treatment cells was collected 72 h post-treatment and cleared by centrifugation at 1,000 x g for 5 min at 4 °C. Cytokine concentrations were quantified using the human Cytokine Antibody Array following the manufacturer’s protocol (#ab133997, Abcam Inc., Toronto, ON, CAN). 1 mL of clarified media was applied per cytokine membrane, followed by an overnight incubation at 4 °C. Chemiluminescence was detected on Blu-Lite films (Dutscher, Bernolsheim, FR) and developed in a dark room using an X-ray film processor. Densitometry was done using the Protein Array Analyzer extension of the ImageJ software (NIH). Spot densitometric values were normalized and corrected for positive and negative controls. Densitometric values were further corrected for cell counts.

### Oxygen consumption and extracellular acidification

The oxygen consumption rates (OCR) and extracellular acidification rates (ECAR) were measured using an XFe96 Seahorse extracellular analyzer from Agilent Technologies Inc. (Santa Clara, CA, USA). Seahorse XFe96 cell cultures microplates (Agilent Technologies Inc.) were pre-coated with 50 µg/mL of Poly-D-Lysine (#A3890401, ThermoFisher Scientific, Waltham, MI, USA) in D-PBS (#311-425-CL, Wisent inc., Saint-Jean-Baptiste, QC, CAN). Seahorse XFe96 sensor cartridges (Agilent Technologies Inc.) were calibrated according to the manufacturer’s protocol. Assay media was prepared as follows for 100 mL: 1 g RPM1640 (with Glucose, without L-glutamine and sodium bicarbonate, #260-012 XK, Wisent inc.), 29.2 mg L-glutamine (#AC386032500, Fisher Chemical, Pittsburgh, PA, USA), 100 µL HEPES 1M (#330-050, Wisent Inc.), 100 mL water, and adjusted pH at 7.2. Cells were plated in Poly-D-Lysine-coated microplates at a density of 45,000 cells/well (A-375), 60,000 cells/well (SK-MEL-5), 70,000 cells/well (MeWo), 60,000 cell/well (WM983B), 65,000 cells/well (IGR-1), 65,000 cell/well (COLO 829), and 70,000 cells/well (MeWo Rho-0) in 100 µL of assay media. Default values for measurement steps were selected on the Wave software (Agilent Technologies Inc.). Sequential measurement steps were: 1) baseline (no injection), 2) oligomycin 3 µM (#ab141829, Abcam Inc.,Toronto, ON, CAN), 3) FCCP 0.5 µM (#15218, Cayman Chemical, Ann Arbor, MI, USA), 4) rotenone 1 µM (#ab143145, Abcam Inc.) and antimycin A 1 µM (#ab141904, Abcam Inc.), 5) injection of monensin 20 µM (#M5273, MilliporeSigma, Oakville, ON, CAN). OCR and ECAR were normalized for cell count. Non-mitochondrial respiration was defined as respiration in cells treated with rotenone and antimycin A and was subtracted from all OCR values. Uncoupled respiration was determined after oligomycin injection. Maximal respiration was determined after Carbonyl cyanide-4 (trifluoromethoxy) phenylhydrazone (FCCP) injection. Spare respiratory capacity was defined as maximal respiration minus basal respiration (untreated). Maximal ECAR was determined after treatment with monensin. Glycolytic reserve was defined as maximal ECAR minus basal ECAR (untreated). For the MeWo and MeWo-Rho-0 comparison, basal (untreated) OCR and ECAR in MeWo were set at 100 %.

### RNA-Seq

Total RNA from MeWo and A-375 cells was extracted as detailed in the Gene expression analysis section. RNA was quantified using Qubit (ThermoFisher Scientific, Mississauga, ON, CAN), and quality was assessed with the 2100 Bioanalyzer (Agilent Technologies, Santa Clara, CA, USA). Transcriptome libraries were generated using the KAPA RNA HyperPrep (Roche, Basel, CHE) using a poly-A selection (ThermoFisher Scientific, Mississauga, ON, CAN). Sequencing was performed on the Illumina NextSeq500 (Illumina, San Diego, CA, USA), obtaining around 30 M single-end 75 bp reads per sample. Samples from the MeWo/BI-D1870 experiments were run independently from those from the A-375/LJH685 experiments. Sequences were trimmed for sequencing adapters and low-quality 3’ bases using Trimmomatic version 0.35 ^81^ and aligned to the reference human genome version GRCh38 (gene annotation from Gencode version 37, based on Ensembl 103) using STAR version 2.7.1a ^82^. Gene expressions were obtained as read count directly from STAR and computed using RSEM ^83^ to obtain normalized gene and transcript level expression, in TPM values, for these stranded RNA libraries. DESeq2 version 1.30.1 ^84^ was used to normalize gene read counts. Sample clustering and principal component analysis (PCA) showed clear and unambiguous clustering of DMSO and RSK inhibitor (BI-D1870, LJH685)-treated groups. Using DESeq2, we introduced a second factor into the model to consider the sample pairing effect by using a likelihood-ratio test to compare the full model and a reduced 35 model with the goal of isolating the condition effect. Removing the pairing effect contribution, we obtain a PCA showing less variance in the PC2 component, representing variability between replicates. Thus, all RNA-Seq analyses consider the batch effect of independent experiments.

### Functional classification and gene set enrichment analyses

Threshold values for fold change > 1.2 fold and false-discovery rate (FDR)-adjusted p-values < 0.1 were used to select differentially expressed transcripts (DET) between groups treated with RSK inhibitors (BI-D1870, LJH685) versus control (DMSO). Log2 fold change and adjusted p-values were calculated with DESeq2. The g:Profiler web-based software ^85^ was used for functional classification of DET based on gene sets from the Reactome pathway database ^86^ and using default options. Gene set enrichment analyses (GSEA) were done using the GSEA software ^87,88^ (v. 4.3.0) using normalized read counts (DESeq2). Undetected genes (0 reads) in all samples were not included in analyses. The Hallmarks MSigDB gene set collection was used for GSEA. Visual representation of GSEA gene sets networks was done using the Cytoscape software ^89^ (v. 3.10.1) with the EnrichmentMap Cytoscape application ^90^. The Molecular signature database (MSigDB) was used for GSEA and network visualization. Enrichment map cutoff parameters were P-value = 0.05 and FDR Q-value = 0.1. Isolated groups of nodes and edges could have been relocated without any modification to increase the comprehensibility of networks. Colored regions were created manually to highlight specific groups of linked gene sets. The figures do not include small groups (e.g. 4 nodes or less).

### Analysis of publicly available databases

RNA expression datasets in human tumors and cell lines were obtained from the open-access and cBioPortal ^91,92^. The human cutaneous melanoma transcriptomics dataset (SKCM) shown here is from the TCGA Research Network: http://www.cancer.gov/tcga (PanCancer Atlas). The human melanoma cell line transcriptomic dataset is from the Cancer Cell Line Encyclopedia (CCLE, Broad, 2019) ^93^. Perturbation effects associated with CRISPR-Cas9 (Public 23Q4+Score, Gene effect: Chronos), RNAi (Achilles+DRIVE+Marcotte, Gene effect: DEMETER2), and inhibitors (PRISM Repurposing Primary Screen, Drug sensitivity/effect: Log2FC of cell count) in human melanoma cell lines were obtained from the DepMap Portal: https://depmap.org/portal. Spearman correlation coefficients (*ρ*) between genes of the SKCM dataset were obtained from cBioPortal. Spearman correlation coefficients between perturbation effects (DepMap) and transcript levels (CCLE) were calculated with Prism (GraphPad, San Diego, CA, USA). Gene essentiality was obtained from DepMap. The MitoCarta3.0 ^94^ dataset and associated MitoPathways3.0 were used to identify and classify mitochondrial proteins associated with transcriptomic data. Heat-maps were generated using Morpheus: https://software.broadinstitute.org/morpheus. For hierarchical clustering and dendrograms, the metric was Euclidean distance, the linkage method was average, and the clustering was done on rows and columns.

### Interaction networks

The STRING web-based resource ^95^ was used to generate an interaction network using the 124-transcript signature associated with resistance to PLK1-targeting approaches (CRISPR-Cas9, RNAi, inhibitors). Mapping settings were set to default: full STRING network (the edges indicate functional and physical associations), meaning of edges based on evidence, with a minimum confidence score of 0.400 (111 proteins involved in interactions, 13 non-interacting proteins). Colored regions were created manually to highlight enriched networks. Network statistics were: 541 edges, 8.73 average node degree, 0.497 average local clustering coefficient, PPI enrichment p-value: < 1.0e-16.

### Statistical analyses

Statistical analyses were done using the Prism software 10_v. 10.2.3 (GraphPad, San Diego, CA, USA) or the Microsoft Excel software All data shown represent the combination mean values obtained from at least 3 independent experiments done at different times with different cell passages. Paired statistical analyses refer to the pairing of samples within independent experiments. Unpaired analyses refer to independent experiments that cannot be paired, such as different cell lines that have not been tested simultaneously. We indicate with a single * symbol the p-values obtained by statistical testing and multiple comparisons below a threshold of 0.05.

## Data availability

The raw and processed data derived from RNA-seq analyses as generated in this study were deposited in NCBI GEO (GSE264686).

## Supplemental information

Document S1. Figures S1-S5.

