## Supplementary material for "Mitochondrial Signatures Shape Phenotype Switching and Apoptosis in Response to PLK1 and RSK Inhibitors in Melanoma": Document S1

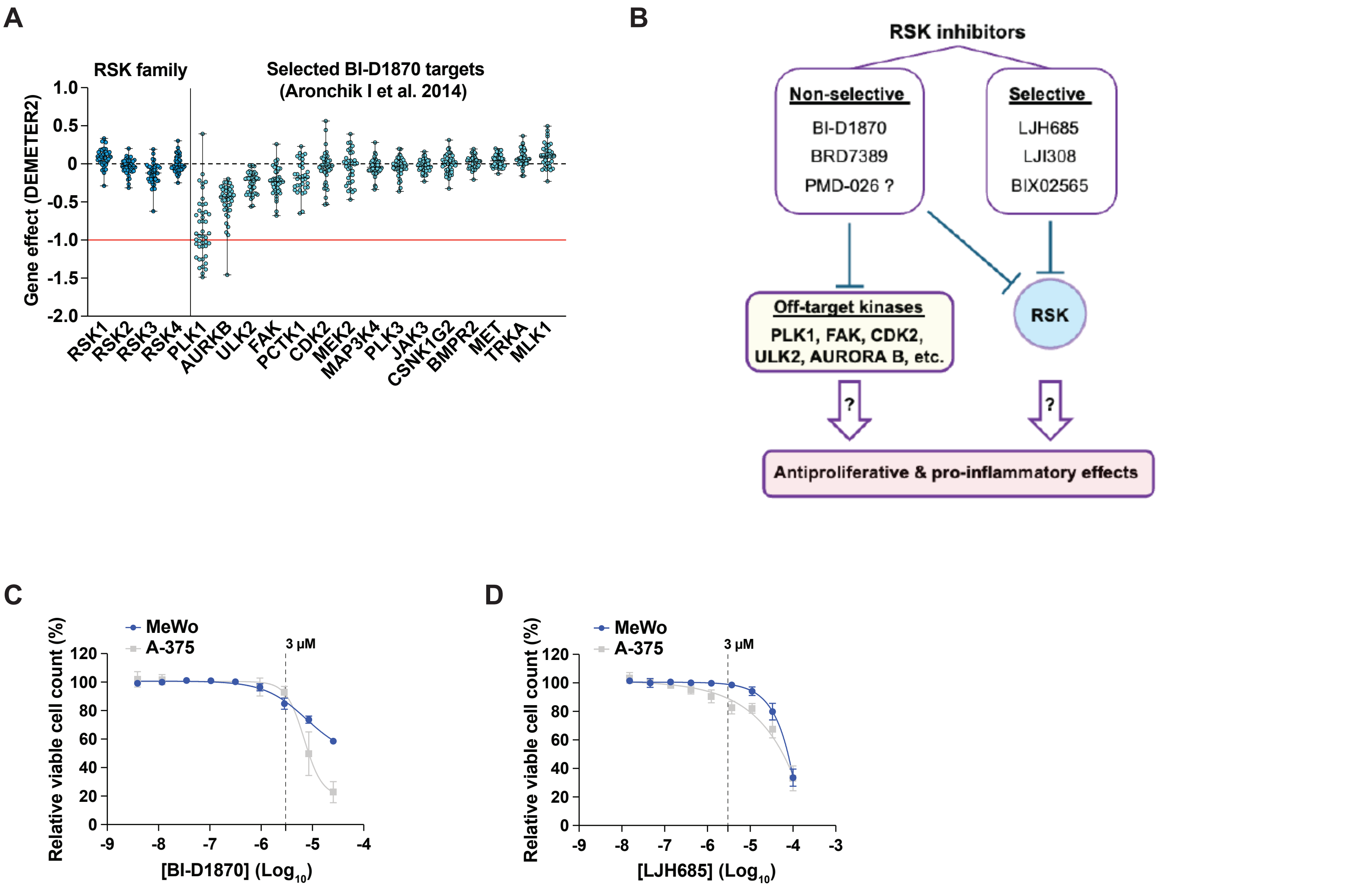


**Figure S1. Impact of RSK inhibitors on proliferation and inflammation.** Related to Figure 1.
(A) Impact of gene silencing (RNAi) of genes shown in Fig. 1A on proliferation dynamics (DEMETER2 algorithm) of human melanoma cell lines (DepMap). The red line (-1) is the median of all common essential gene scores. Each dot represents a single cell line; data are shown as median +/- min/max.
(B) Research hypothesis. Off-target effects of  non-selective RSK inhibitors could mediate their antiproliferative and pro-inflammatory effects.
(C,D) Relative viable cell counts of MeWo and A-375 cells treated with serial dilutions of RSK inhibitors BI-D1870 and LJH685 for 72 h. The dotted line indicates impact on cell growth at a dose of 3 µM. Data are shown as mean +/- SEM from 3 independent experiments. Related to Figure 1C, 1D and 1E.

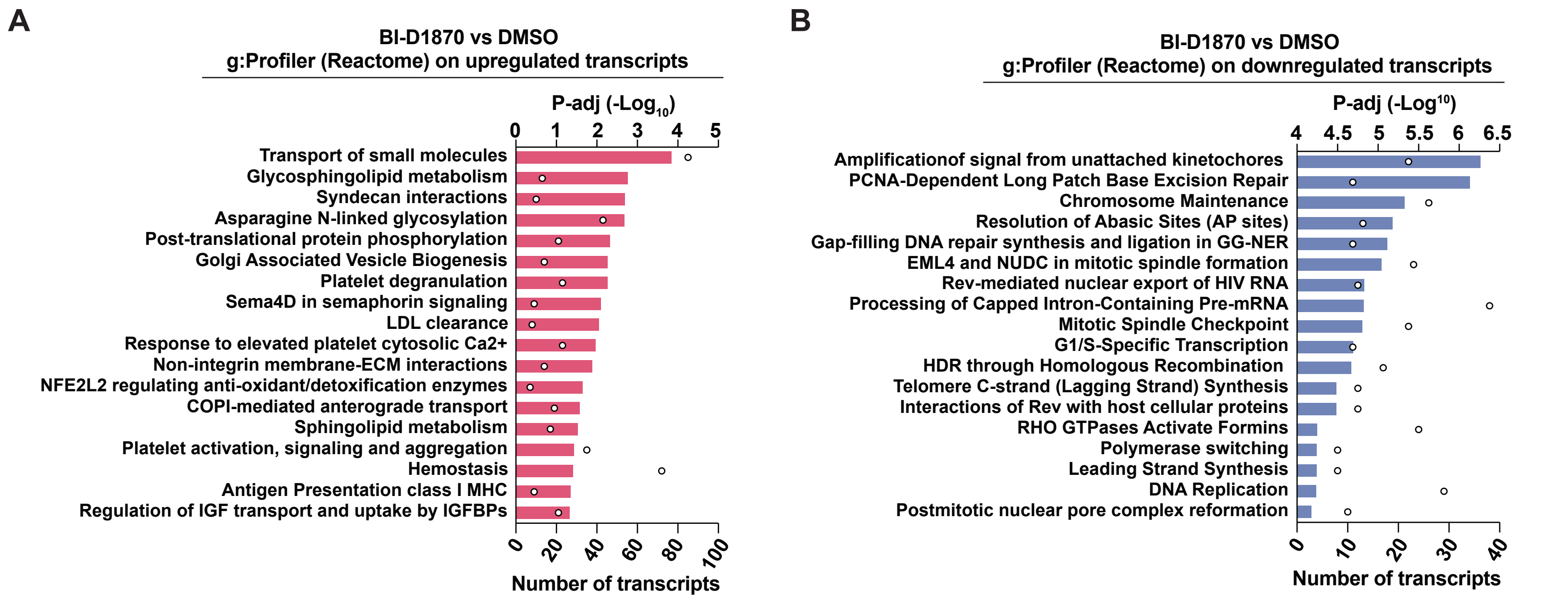


**C**

**GSEA : hallmarks top 9 (BI-D1870 in MeWo)**

| Name | ES | NES | NOM p | FDR q |
|---|---|---|---|---|
| EMT | 0.48873648 | 2.5246518 | 0 | 0 |
| Inflammatory response | 0.45301688 | 2.2891004 | 0 | 0 |
| UV response DN | 0.4419671 | 2.1537316 | 0 | 0 |
| IL6 JAK STAT3 signaling | 0.47230023 | 2.0519352 | 0 | 4.10E-04 |
| Protein secretion | 0.4386726 | 1.9959487 | 0 | 6.43E-04 |
| TNFA signaling via NFKB | 0.38886443 | 1.9912329 | 0 | 5.36E-04 |
| Apical junction | 0.38008124 | 1.9361606 | 0 | 4.60E-04 |
| TGF Beta signaling | 0.4807785 | 1.9128569 | 0 | 6.35E-04 |
| Heme metabolism | 0.37420398 | 1.8895833 | 0 | 7.33E-04 |
| E2F targets | -0.5721137 | -2.8329363 | 0 | 0 |
| G2M checkpoint | -0.47602367 | -2.3789809 | 0 | 0 |
| MYC targets V2 | -0.57238406 | -2.2582335 | 0 | 0 |
| MYC targets V1 | -0.4141819 | -2.0644515 | 0 | 0 |
| Pancreas beta cells | -0.49898466 | -1.6700398 | 0.017077798 | 0.004277723 |
| DNA repair | -0.30767232 | -1.467096 | 0.019400353 | 0.025454987 |
| WNT beta catenin signaling | -0.32602665 | -1.1845939 | 0.21578947 | 0.2025009 |
| Spermatogenesis | -0.19413668 | -0.8766633 | 0.73193914 | 0.8223576 |
| Notch signaling | -0.246087 | -0.8713314 | 0.6386861 | 0.7431215 |

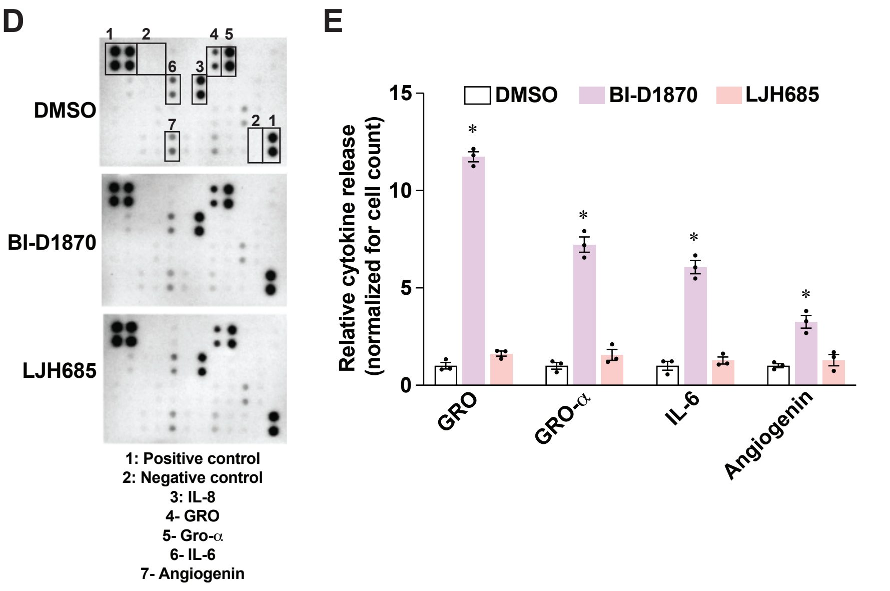


**Figure S2. Characterization of the pro-inflammatory response of BI-D1870 in MeWo cells.** Related to Figure 2.
(A) Gene functional analysis (g:Profiler; Reactome) of additionnal significantly upregulated transcripts, related to Figures 2B and 2C.
(B) Gene functional analysis (g:Profiler; Reactome) of additionnal significantly downregulated transcripts, related to Figures 2B and 2D.
(C) Top gene set enrichments identified by GSEA (Hallmarks, MSigDB) in MeWo cells treated with BI-D1870 (3 μM) for 72 h versus DMSO controls. ES: enrichment score; NES: normalized enrichment score; NOM p: nominal p-value; FDR q: false discovery rate q-value. Related to Figure 2E.
(D) Representative films of cytokine arrays and HRP chemiluminescence associated with the presence of cytokines in conditionned media from MeWo cultures. Related to Figures 2H and 2I.
(E) Densitometric analysis of selected cytokines shown in D. Data are shown as means of 3 independent experiments +/- SEM. * $p < 0.05$, One-Way ANOVA with Dunnett multiple comparison tests.

**A**

**GSEA : hallmarks top 8 (LJH685 in A-375)**

| Name | ES | NES | NOM p | FDR q |
|---|---|---|---|---|
| WNT beta catenin signaling | 0.44859722 | 1.6167767 | 0.00660793 | 0.026256047 |
| Coagulation | 0.30320382 | 1.3754022 | 0.02970297 | 0.12270627 |
| Hedgehog signaling | 0.3405547 | 1.3754022 | 0.19058824 | 0.40981713 |
| Spermatogenesis | 0.20565586 | 0.89566976 | 0.6876404 | 1 |
| Adipogenesis | 0.1780325 | 0.8552081 | 0.83589745 | 1 |
| Notch signaling | 0.23504652 | 0.77731514 | 0.8242009 | 1 |
| MYC targets V2 | 0.14803095 | 0.571583 | 0.991342 | 1 |
| G2M checkpoint | 0.09866795 | 0.46900535 | 1 | 0.9999545 |
| Hypoxia | -0.46831548 | -2.1386015 | 0 | 0.001448276 |
| Inflammatory response | -0.4711206 | -2.129727 | 0 | 7.24E-04 |
| TNFA signaling via NFKB | -0.46115926 | -2.1272213 | 0 | 4.83E-04 |
| KRAS signaling up | -0.46687463 | -2.121482 | 0 | 3.62E-04 |
| IL2 STAT5 signaling | -0.44417548 | -2.0482023 | 0 | 2.9E-04 |
| P53 pathway | -0.43288162 | -1.97651 | 0 | 2.41E-04 |
| Estrogen response late | -0.43226212 | -1.9640619 | 0 | 2.07E-04 |
| Xenobiotic metabolism | -0.4247412 | -1.9147873 | 0 | 3.83E-04 |

**B**

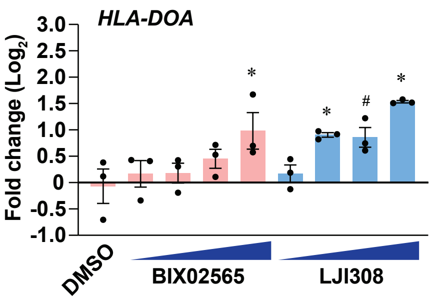


**Figure S3. Characterization of immune effects of LJH685 in A-375 cells.** Related to Figure 3.
(A) Top gene set enrichments identified by GSEA (Hallmarks, MSigDB) in A-375 cells treated with LJH685 (3 µM) for 72 h versus DMSO controls. ES: enrichment score; NES: normalized enrichment score; NOM p: nominal p-value; FDR q: false discovery rate q-value. Related to Figure 3F.
(B) *HLA-DOA* transcript levels in A-375 cells treated with RSK inhibitors for 72 h. Doses: 0.3, 1, 3, and 10 µM. Data shown as mean +/- SEM from 3 independent experiments. * $p < 0.05$, # $p < 0.1$, One-Way ANOVA with Dunnett multiple comparison test. Related to Figures 3I and 3J.

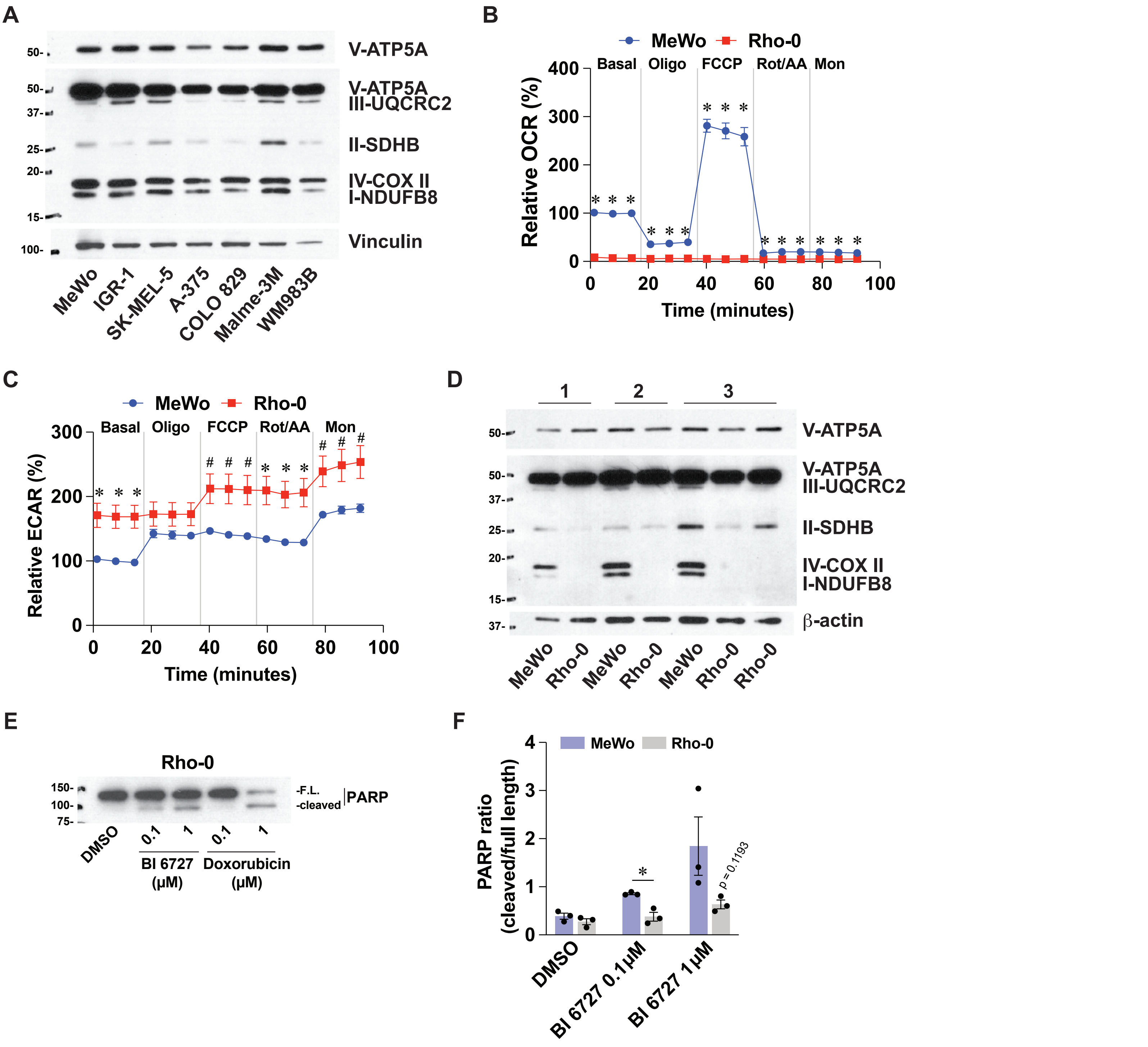


**Figure S4. Metabolic characterization of melanoma cell lines and Rho-0 model.** Related to Figure 6.
(A) Immunoblotting of OXPHOS complex I-V proteins in protein extracts from selected melanoma cell lines. Representative of 3 independent experiments.
(B) Oxygen consumption rate (OCR) in Rho-0 cells and parental controls (MeWo). Data is relative to MeWo basal conditions. Details about acute treatments in the Materials and methods section.
(C) Extracellular acidification rate (ECAR) in Rho-0 cells and parental controls (MeWo), as indicated in B.
(D) Immunoblotting of OXPHOS complex I-V proteins in Rho-0 cells and parental controls (MeWo). Numbers refer to independent experiments.
(E) Immunoblotting for PARP in protein extracts from MeWo-Rho-0 cells treated with DMSO 0.1 %(v/v), BI 6727, or doxorubicin for 72 h. F.L: full length.
(F) Densitometric analysis of 3 independent experiments related to E.
Data are shown as mean +/- SEM from independent experiments. * $p < 0.05$, # $p < 0.10$, paired Student's *t*-tests.

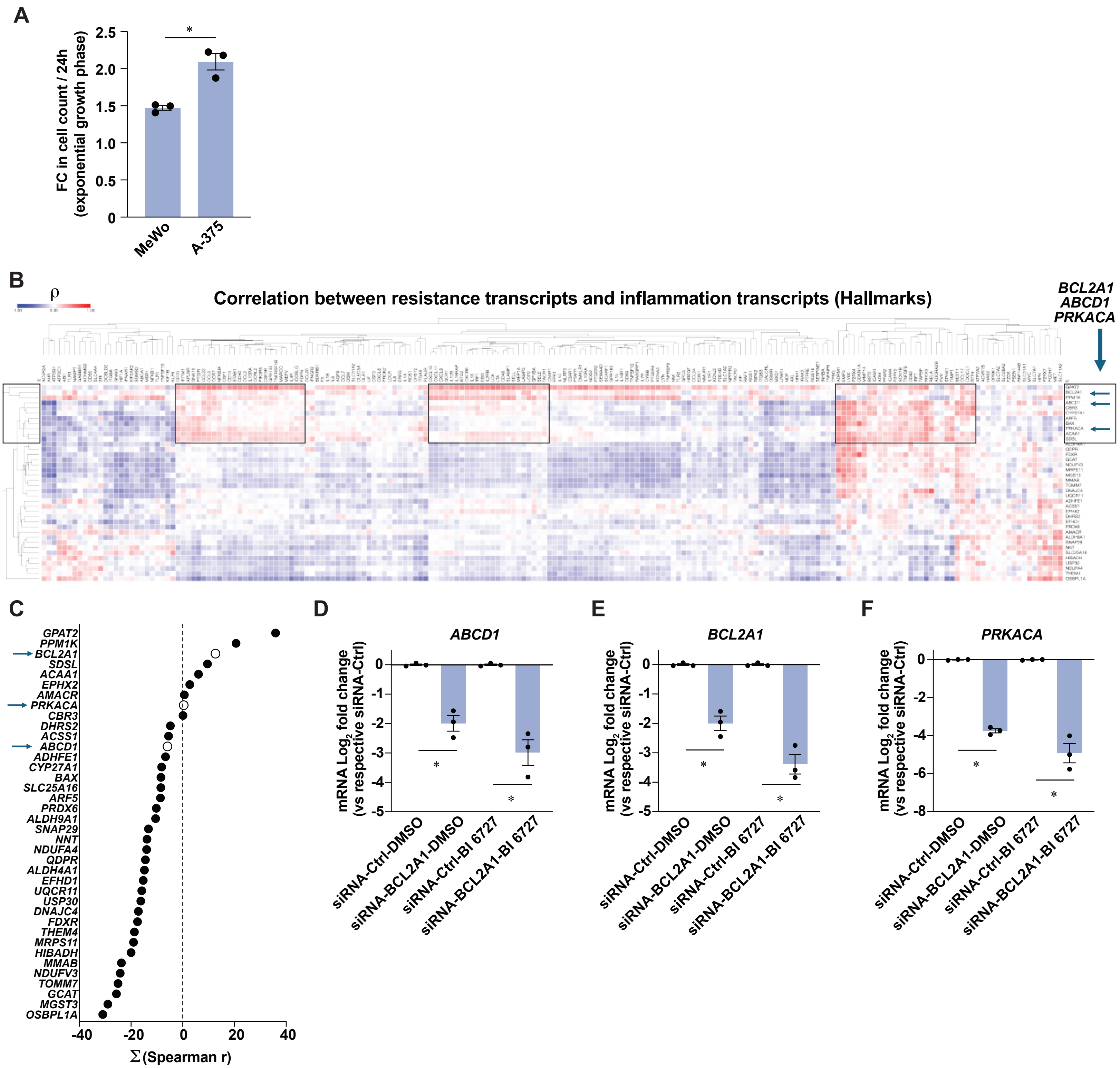


**Figure S5. Characterization of melanoma cell lines and identification of mitochondrial targets.** Related to Figure 7.
(A) Fold change in cell count during the exponential growth phase of MeWo and A-375 cells.
(B) Heat-map of Spearman correlation coefficients (ρ) between PLK1-targeting resistance transcripts (vertical) and transcripts from Inflammatory Response gene list (Hallmarks, MSigDB), using RNA-Seq data from skin cutaneous melanoma tumors (TCGA, PanCancer Atlas). Boxes indicates clusters associated with inflammation. Arrows indicate tested genes in this study.
(C) Summation of all Spearman correlations from B, for each transcript encoding a mitochondrial protein associated with resistance to PLK1 targeting. Arrows indicate tested genes in this study.
(D-F) Expression levels of *ABCD1*, *BCL2A1*, and *PRKACA* in MeWo cells transfected with non-targeting siRNA-Ctrl or siRNA pools against selected genes for 48 h and treated with DMSO 0.1 % (v/v) or BI 6727 (100 nM) for 72 h. Data are shown as mean +/- SEM from 3 independent experiments. * $p < 0.05$, paired (C-E) and unpaired (A) Student's *t*-tests.
